# Doublecortin restricts neuronal branching by regulating tubulin polyglutamylation

**DOI:** 10.1101/2023.06.02.543327

**Authors:** Muriel Sébastien, Alexandra L. Paquette, Emily N. P. Prowse, Adam G. Hendricks, Gary J. Brouhard

## Abstract

Doublecortin (DCX) is a neuronal microtubule-associated protein (MAP) that binds directly to microtubules via two Doublecortin (DC) domains. The DC domains sense the nucleotide state, longitudinal curvature, and protofilament number of the microtubule lattice, indicating a role in the regulation of microtubule structure in neurons. Mutations in DCX cause lissencephaly and subcortical band heterotopia (also known as double-cortex syndrome) due to impaired neuronal migration. To better understand the role of DCX in neuronal migration, we developed a model system based on induced pluripotent stem cells (iPSCs). We used CRISPR/Cas9 to knock-out the *Dcx* gene in iPSCs and differentiated the cells into cortical neurons. Compared to control neurons, the DCX-KO neurons showed reduced velocities of nuclear movements. The reduced velocities coincided with an increase in the number of neurites early in neuronal development, consistent with a neuronal migration phenotype and previous findings in a DCX-KO mouse model. Neurite branching is regulated by a host of MAPs and other factors, as well as by microtubule polymerization dynamics. However, EB comet dynamics were unchanged in DCX-KO neurons, with similar growth rates, lifetimes, and numbers. Rather, we observed a significant reduction in α-tubulin polyglutamylation in DCX-KO neurons. Polyglutamylation levels and neuronal branching were rescued by expression of DCX or of TTLL11, an α-tubulin glutamylase. Using U2OS cells as an orthogonal model system, we show that DCX and TTLL11 act synergistically to promote polyglutamylation. Polyglutamylation regulates numerous MAPs, severing enzymes, and molecular motors. Consistently, we observe that lysosomes in DCX-KO neurons show a reduction of their processivity. We propose that the DCX acts as a positive regulator of α-tubulin polyglutamylation and restricts neurite branching. Our results indicate an unexpected role for DCX in the homeostasis of the tubulin code.

## Introduction

During human brain development, approximately 10^10^ newborn neurons migrate out of their stem cell niche to form the cerebral cortex (Herculano-Houzel 2009). Migrating neurons generally have two slender processes that extend from the cell body (Rakic 1972). Their migration can be divided into two steps: (1) elongation of the leading process and (2) translocation of the cell body in the direction of motion (Nadarajah and Parnavelas 2002). Both steps rely heavily on the neuronal microtubule cytoskeleton (Kapitein and Hoogenraad 2015). In the leading process, microtubules contribute pushing forces (Dogterom et al. 2005), activate and reinforce actin polymerization, respond to guidance cues, and steer neuronal migration (Menon and Gupton 2016). In the cell body, microtubules and motor proteins generate forces that pull the nucleus forward (Umeshima, Hirano, and Kengaku 2007). In addition, microtubules serve as roadways for the transport of organelles, synaptic vesicle precursors, and neurotrophic factors by motor proteins (Maday et al. 2014).

To enable these diverse functions, the microtubule network is regulated and patterned at multiple levels. First, in neurites, microtubules are dynamic in neurite tips and stable in neurite shafts (Baas and Black 1990). In axons, microtubules have uniform orientation (plus-end out); in dendrites, microtubule orientation is mixed (Baas et al. 1988). Second, microtubules are altered by post-translational modifications (PTMs), known as the “tubulin code” (Verhey and Gaertig 2007; Magiera et al. 2018; Roll-Mecak 2020), to Sébastien et al. 2024 (preprint) create subpopulations of microtubules with variable chemical and mechanical properties (Magiera et al. 2018; Roll-Mecak 2020). Third, the tubulin code governs a host of microtubule-associated proteins (MAPs), including structural MAPs (Ramkumar, Jong, and Ori-McKenney 2018; Bodakuntla et al. 2019), polymerases and depolymerases (Howard and Hyman 2007), plus-end and minus-end regulators (Akhmanova and Steinmetz 2015), and motor proteins (Hirokawa et al. 2009; Verhey and Ohi 2023). Each MAP has a specific localization, affinity, and function towards microtubules. Together, these regulatory mechanisms create complex microtubule “cartography” (Iwanski and Kapitein 2023). For example, dendritic microtubules are organized into adjacent bundles that differ in their polarity and their PTMs (Tas et al. 2017). Dysregulation of MAPs or loss of microtubule patterning can lead to brain malformation, neurodegeneration, and/or impaired neural function.

Human brain formation depends critically on the MAP Doublecortin (DCX) (Francis et al. 1999; Gleeson et al. 1999). The *Dcx* gene is located on the X-chromosome and was discovered in 1998 in a cohort of patients with double cortex syndrome and X-linked lissencephaly (Gleeson et al. 1998; des Portes et al. 1998). These diseases are caused by mutations in DCX that lead to a failure in the radial migration of cortical neurons and result in a disorganized lamination of the cerebral cortex (Feng and Walsh 2001). How DCX mutations cause migration defects remains only partially understood, in part due to competing hypotheses about the structural role of DCX in microtubule regulation.

Most disease-causing mutations in DCX are missense mutations that cluster in a pair of tandem globular domains, known as the DC domains (Sapir et al. 2000; Taylor et al. 2000). DC domains bind to microtubules at the vertex of 4 tubulin dimers (Moores et al. 2004; Fourniol et al. 2010), a binding site shared with end-binding (EB) family proteins (Maurer et al. 2012) and doublecortin-like kinase 1 (DCLK1)(Omori et al. 1998; Kim et al. 2003). The location of this binding site, in close proximity to inter- and intra-protofilament interfaces, makes DC domains (and EBs) highly responsive to the structural state of the microtubule lattice.

The N-terminal domain, DC1, is tightly folded (Kim et al. 2003) and binds to straight GDP lattices (Fourniol et al. 2010). The second domain, DC2, is only partly folded (Kim et al. 2003; Burger et al. 2016) and binds GDP-Pi lattices (Manka and Moores 2018), curved lattices (Bechstedt, Lu, and Brouhard 2014), and other tubulin intermediates (Moores et al. 2006; Manka and Moores 2020). Together, the DC domains enable DCX to bind cooperatively to the microtubule lattice (Bechstedt and Brouhard 2012; Rafiei et al. 2022). In addition, the DC domains bind in close proximity to kinesin motor domains on microtubules (Liu et al. 2012) and impact kinesin-1 and dynein motility *in vitro* (Monroy et al. 2020; Fu et al. 2022). Many of these diverse structural interactions are disrupted by patient mutations in the DC domains (Bechstedt and Brouhard 2012; Bechstedt, Lu, and Brouhard 2014; Manka and Moores 2020), suggesting a direct role for DCX in regulating microtubule structure and intracellular trafficking. Yet how mutations in DC domains disrupt function in the full context of the neuronal proteome remains unclear, especially at neurite tips and cell bodies where DCX is enriched (Friocourt et al. 2003; Tint et al. 2009; Dema et al. 2024).

The link between the disease phenotype seen in patients and the mechanistic behavior seen in reconstitution assays has been partially elucidated by several *in vitro* and *in vivo* models. Surprisingly, *Dcx*^*-/y*^ (hemizygote) mice have very mild phenotypes (Corbo et al. 2002), with a cortical lamination indistinguishable from wildtype; only a double knock-out of DCX and DCLK1 gives rise to a disorganized cortex (Deuel et al. 2006). Stronger phenotypes were observed with shRNA-mediated knock-down of DCX in rat embryos, which resulted in severe cortical migration defects (Bai et al. 2003). This RNAi approach was shown to have significant off-target effects mediated by the miRNA *let7* (Baek et al. 2014), complicating the interpretation of shRNA-mediated knock-down experiments (Liu et al. 2012). However, explants of neurons from *Dcx*^*-/y*^ mice allowed for an initial characterization of cortical neuron behavior: DCX knock-out disrupted the movement of the nucleus (nucleokinesis) and increased neurite branching (Kappeler et al. 2006; Koizumi et al. 2006). So far, these phenotypes have not been linked to a clear mechanism of microtubule regulation. Basic cellular data on microtubules remains unmeasured, such as whether microtubule dynamics and intracellular trafficking are perturbed in non-shRNA-mediated neuronal models.

To better understand DCX’s role in neuronal migration, we took advantage of human induced pluripotent stem cells (iPSCs) and CRISPR/Cas9 gene editing. Human iPSC models have the following advantages: (1) human-specific genotypes, which are significant in the context of cortical development (Buchsbaum and Cappello 2019); (2) relatively simple knock-in of patient mutations in an isogenic background; and (3) direct observation of neurons immediately after terminal differentiation as opposed to harvesting and plating of neurons from E18 rodent brains. Thus, we generated a series of DCX-KO and patient mutant iPSC models.

In our iPSC models, DCX-KO neurons showed impaired nucleokinesis and increased neurite branching, consistent with phenotypes in *Dcx*^*-/y*^ rodent studies (Kappeler et al. 2006; Koizumi et al. 2006). Using our DCX-KO model, we find that excessive branching is the result of higher rates of branch initiation, a pathway known to involve severing enzymes and structural MAPs. In addition, the processivity of lysosomes is impaired in DCX-KO neurons, linking DCX directly to intracellular trafficking by kinesins and dynein. These diverse phenotypes involve multiple protein networks, suggesting disruption of an upstream regulator of the microtubule network. While searching for a common mechanism, we made the unexpected observation that DCXKO neurons exhibit a significant reduction in α-tubulin polyglutamylation, one of the central PTMs of the neuronal tubulin code. Expressing exogenous DCX in DCX-KO neurons was sufficient to rescue polyglutamylation levels and to restrict neurite branching back to control. Rescue was also achieved by over-expression of TTLL11, an α-tubulin specific polyglutamylase, or by driving polyglutamylation with paclitaxel. Using U2OS cells as an alternate model, we established that co-expression of DCX and TTLL11 was sufficient to create a significant, synergistic increase in polyglutamylation. Finally, we confirmed that polyglutamylation levels are reduced in 6 patient mutant knock-in neuronal models, linking our findings on the tubulin code to patient genotypes. Our results on DCX and polyglutamylation demonstrate that MAPs not only “read” the tubulin code but also regulate it.

## Results

### Nuclear mobility is impaired in DCX-KO cortical neurons

To understand the role of DCX in regulating the neuronal cytoskeleton, we used CRISPR/Cas9 genome editing to knock-out the expression of DCX in a male iPSC line. We designed a gRNA to target a sequence upstream of the first microtubule binding domain DC1, resulting in multiple iPSC clones expressing only N-terminal fragments (Fig. 1A and Fig. S1A-B). We confirmed DCX knock-out by Western blots and immunohistochemistry using antibodies against DCX’s C-terminal domain (Fig. 1B-C and Fig. S1C). The morphology of our DCX-KO cells was indistinguishable from control cells at the iPSC stage as well as after induction to neural progenitor cells (NPCs, data not shown). Terminal differentiation of DCX-KO NPCs into cortical neurons was confirmed by immunohistochemistry and Western blots against β3-tubulin (Tubb3), a neuron-specific isotype (Fig. 1C and Fig. S1D). At DIV3 (3 days in terminal differentiation medium), iPSC-derived cortical neurons showed a bipolar morphology reminiscent of migrating neurons, with two or more slender processes that extended from the cell body (Rakic 1972; Nadarajah and Parnavelas 2002).

**Figure 1:**
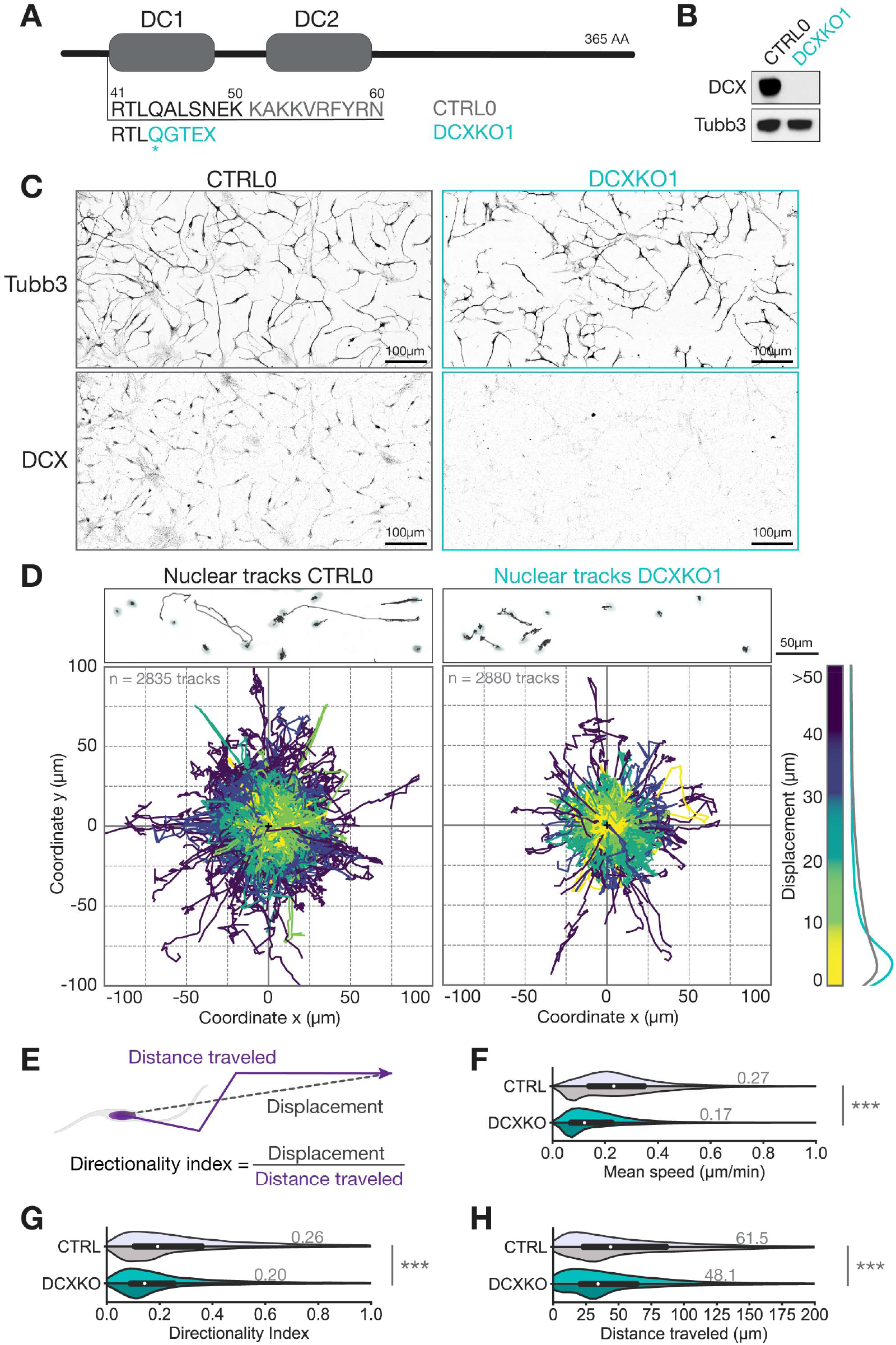
Nuclear mobility is impaired in DCX-KO cortical neurons. **(A)** DCX contains two microtubule binding domains, DC1 and DC2. DCX-KO cells were obtained by indels introduced in *Dcx*’s gene sequence before DC1. CTRL0 and DCXKO1 amino acid sequences are shown from residue 41 to 60. The asterisk shows the indel location and the resulting amino acid sequence in DCXKO1 is shown in cyan. **(B)** Immunoblots of CTRL0 and DCXKO1 neuron lysates at three days of differentiation (DIV3), with antibodies against Tubb3 and DCX. The same samples were loaded on different gels for both antibodies. **(D)** (top panel) Still images of live neurons with Syto nuclear dye. Nuclei that are detected by trackmate are circled in light blue, and their trajectories over the 8h period of imaging are shown in grey. (Bottom panel) Position plots showing a sample of trajectories extracted from CTRL0 (left) and DCXKO1 (right) nuclei tracking. All nuclei are considered to be at the same coordinates (0,0) at their first timepoint. Tracks are color-coded according to the displacement (distance between the first and the last position) of the nucleus. Distribution of displacement for both genotypes are shown on the right end side of the panel. *n*=2835 and 2880 nuclei respectively for CTRL0 and DCXKO1. **(E)** Schematics showing the displacement, distance traveled, and directionality index parameters. **(F)** Distribution of mean speed achieved by CTRL and DCXKO nuclei during the imaging period. **(G)** Distribution of directionality index or efficiency of the trajectory for CTRL and DCXKO nuclei. Higher directionality index indicates more unidirectional movement. **(H)** Distribution of distance traveled by CTRL and DCXKO nuclei during the imaging period. For F-H) Violin plots are split between the two control (CTRL0, light grey and CTRL1, dark grey) and two DCXKO (DCXKO1, cyan and DCXKO4, dark cyan) lines used in this study. White dots are showing median values, while the mean values are detailed on each plot. The thick black lines represent the quartiles. These three plots were cut-off to better show the main population, but the maximum values for distance traveled, mean speed and directionality index are 497 µm, 2.7 µm/min and 1 respectively. *n*=21826 tracks for CTRL0, 24801 tracks for CTRL1, 6461 tracks for DCXKO1 and 14837 tracks for DCXKO4. Each line was used in 3 paired experiments, tracks with less than 30min of imaging time were excluded from the dataset. Statistical analysis was performed using the Kruskal-Wallis non parametric H test: *** *p*<0.001.

To determine if DCX-KO cortical neurons show phenotypes consistent with lissencephaly, we first measured the translocation of nuclei in control and DCX-KO cells using automated tracking algorithms (Fig. 1D-H, Fig. S2, and Videos S1-S2). Representative images of nuclei and their tracks are shown in Figure 1D (top panel), above 2-dimension plots of nuclear position (n ≅ 2800 tracks) over an 8 h observation window. These tracks showed a broad distribution of nuclear displacements (distance between first and last coordinates, Fig. 1E) in all directions, as some nuclei move significantly more than others of the same genotype. Despite this heterogeneity, DCX-KO neurons have a greater proportion of small displacements (<10 µm) while control neurons have more large displacements (>10 µm) (Fig. 1D, right panel and Fig. S2A, right panel). To mitigate the effect of false-positive tracks from our algorithm, only tracks that stayed visible for ≥6 frames (30 min) through the whole imaging period were analyzed (Fig. 1F-H and Fig. S2B-C). The nuclear movements were also slower. In control neurons, the nuclei had a mean speed of 0.27 ± 0.01 µm/min on average over the observation window (Fig. 1F). In contrast, in DCX-KO neurons, the nuclei had a significantly reduced speed of 0.17 ± 0.01 µm/min on average over the same window (*p*<0.0001, Fig. 1F). Moreover, we measured the maximum speed, or the fastest segment of each nucleus track (Fig. S2B). DCX-KO nuclei had reduced maximum speeds compared to control nuclei, suggesting a generalized decrease in speed as opposed to increased pausing. In parallel, we measured a “directionality index”, which is the ratio between displacement and distance traveled, or the efficiency of the movement (Fig. 1E and 1G). A perfectly straight trajectory would result in a directionality index of 1. The directionality index for both genotypes are relatively low as the direction of movement is random and not chemotactic or durotactic. However, DCX-KO neurons have a reduced directionality index compared to control (DCX-KO 0.20 ± 0.01 vs control 0.26 ± 0.01, *p*<0.0001), implying that they are 20% less efficient in their displacement relative to control neurons (Fig. 1G). This inefficiency causes DCX-KO nuclei to travel shorter distances (Fig. 1H, control 61.5 ± 0.2 µm vs DCX-KO 48.1 ± 0.3 µm, *p* <0.0001). We conclude that DCX-KO nuclei are slower, cover less distance, and are less directional, consistent with the nucleokinesis defects observed in *Dcx*^*-/y*^ explant neurons (Kappeler et al. 2006; Koizumi et al. 2006; Belvindrah et al. 2017).

### DCX-KO neurons establish more branches through increased rates of initiation and retraction

Nucleokinesis is one of two steps in the process of neuronal migration; the other step being neurite extension (Dogterom and Koenderink 2019; Tsai and Gleeson 2005; Coles and Bradke 2015). As expected, DCX is enriched at neurite tips in our cell model (Fig. S4E), so we wondered if a defect in this compartment could also contribute to defects in neuronal migration. To test this idea, we imaged individual neurites using either phase contrast microscopy or fluorescence microscopy with live cell fluorescent labels for microtubules and actin filaments (SiR-Tubulin and CellMask Actin, respectively). As shown in Fig. 2A-C (Videos S3-S6), we observed the canonical sequence of events (Kalil and Dent 2014; Pacheco and Gallo 2016): (A) first, neurites were initiated when a transient burst of actin polymerization was stabilized by microtubules; (B) the initiated neurite elongated at a relatively consistent velocity; (C) in most cases, the neurite later retracted. The balance of initiation and retraction events determines the net increase in neurite number per cell. We measured no significant change between control and DCX-KO cells in the rates of elongation or shrinkage (Fig. 2D-E); rather, only initiation and retraction events were affected. In DCX-KO neurons, the number of initiation events increased (Fig. 2G), as did the number of retraction events, albeit to a lesser extent (Fig. 2F), resulting in an imbalance relative to controls. We measured a 2-fold increase in neurite formation (number of neurite initiations minus number of neurite retractions) per cell in DCX-KO neurons (Fig. 2H). We confirmed that DCX-KO increases neurite number using both types of live-cell imaging as well as direct counting of neurites in immunocytochemistry of fixed neurons (Fig. 2H and 2J). Figures 2D-H report the results from phase contrast microscopy of unlabeled cells to avoid any confounding effects of the fluorescent reagents. To account for variations in cell developmental stage, we also measured the neurite density, or the number of neurites divided by the total length of the cell’s neurites (Fig. 2I and Fig. S3) and found again a significant increase in DCX-KO neurons. We conclude that DCX plays a significant role in restricting neurite formation (Kappeler et al. 2006; Bilimoria et al. 2010; Li et al. 2014). Cortical neurons with an excessive number of neurites may struggle to migrate, because each branch pulls the cell body in a different direction (Martini et al. 2009; Chai et al. 2015; Gupta et al. 2003). These results are therefore consistent with hypotheses that excessive branching may contribute to the neuronal migration defects observed in patients with lissencephaly and double cortex syndrome (Koizumi et al. 2006).

**Figure 2:**
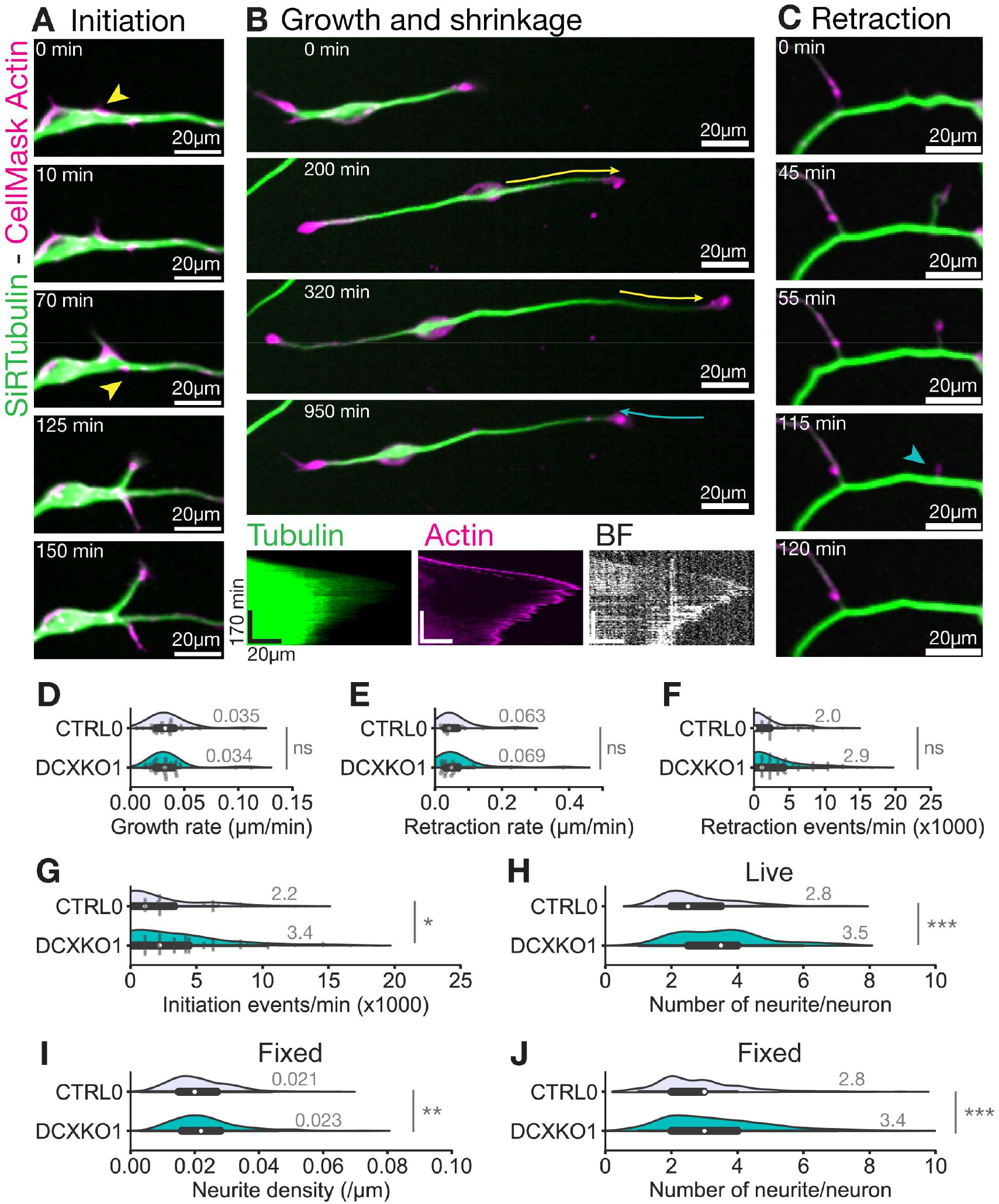
DCX-KO neurons establish more branches through increased rates of initiation/retraction. **(A) (B)** and **(C)** Representative images of a neurite initiation (A), growth/shrinkage (B) and retraction (C) events in a CTRL0 neuron recorded after incubation with tubulin and actin binding dyes: SiR-tubulin (green) and Cellmask actin (magenta). Yellow arrowheads show actin protrusions where a new neurite is initiated. cyan arrowheads show the remains of a neurite retracting completely. Yellow arrows show growth event and cyan arrows show retraction events. These events were used for calculation of growth and retraction speed (D and E), by measuring neurite length and lifetime on kymographs. An example kymograph is shown in the bottom panel in B with brightfield and fluorescence imaging. **(D)** Distribution of neurite growth rate in CTRL0 and DCXKO1 neurons. **(E)** Distribution of neurite retraction rate in CTRL0 and DCXKO1 neurons. For D-E, *n*=26 control and 38 DCX-KO events, from 2 independent experiments. Statistical analysis was performed using a Mann-Whitney U test: ns, nonsignificant. **(F)** Distribution of the number of neurite retraction events/min seen in CTRL0 and DCXKO1 neurons. Cells showing no retraction events were included in the analysis, but the points were not represented on the plot. **(G)** Distribution of the number of neurite initiation events/minute per neuron in CTRL0 and DCXKO1 neurons. Cells showing no initiation events were included in the analysis, but the points were not represented on the plot. **(H)** Number of neurites found per neuron in CTRL0 and DCXKO1 neurons. For F-H, *n*=68 control and 93 DCX-KO neurons from 2 independent experiments. Statistical analysis was performed using a Mann-Whitney U test: ns nonsignificant, * *p*<0.05, *** *p*<0.001. **(I)** Violin plot showing the distribution of neurite density per neuron for CTRL0 and DCXKO1 neurons in fixed samples. **(J)** Violin plot showing the distribution of neurite number per neuron for CTRL0 and DCXKO1 neurons in fixed samples. For I-J, *n*=323 control and 351 DCX-KO neurons from 3 independent experiments. Statistical analysis was performed using a Mann-Whitney U test: ** *p*< 0.01, *** *p*<0.001.

### Microtubule dynamics are unaffected in DCX-KO neurons

What could cause the increase in neurite branching in DCX-KO neurons? Microtubule dynamics play a major role in branching via the polymerization of microtubules into nascent branches (Dehmelt and Halpain 2004; Kawauchi and Hoshino 2008; Cooper 2013). Hence, one plausible explanation for the branching phenotype is altered microtubule dynamics, whereby increased microtubule lifetimes/numbers in DCX-KO neurons stabilize a greater percentage of the transient bursts of actin polymerization. In reconstitution assays, DCX promotes microtubule nucleation (Moores et al. 2004; Bechstedt and Brouhard 2012), reduces the microtubule catastrophe frequency (Manka and Moores 2020), and slows their post-catastrophe shrinkage (Moores et al. 2006). But DCX appears to be excluded from microtubule plus ends by end-binding proteins in cell lines (Ettinger et al. 2016), so the potential impact of DCX-KO on microtubule dynamics in neurons is difficult to predict. As a first approach to look for changes in the microtubule cytoskeleton in DCX-KO neurons, we stained for α-tubulin and the neuronal specific Tubb3, as markers of the total microtubule population (Fig. 3A and S4A-B). We found that DCX-KO neurons have equivalent levels of total tubulin, albeit with a slight reduction in the intensity of Tubb3.

**Figure 3:**
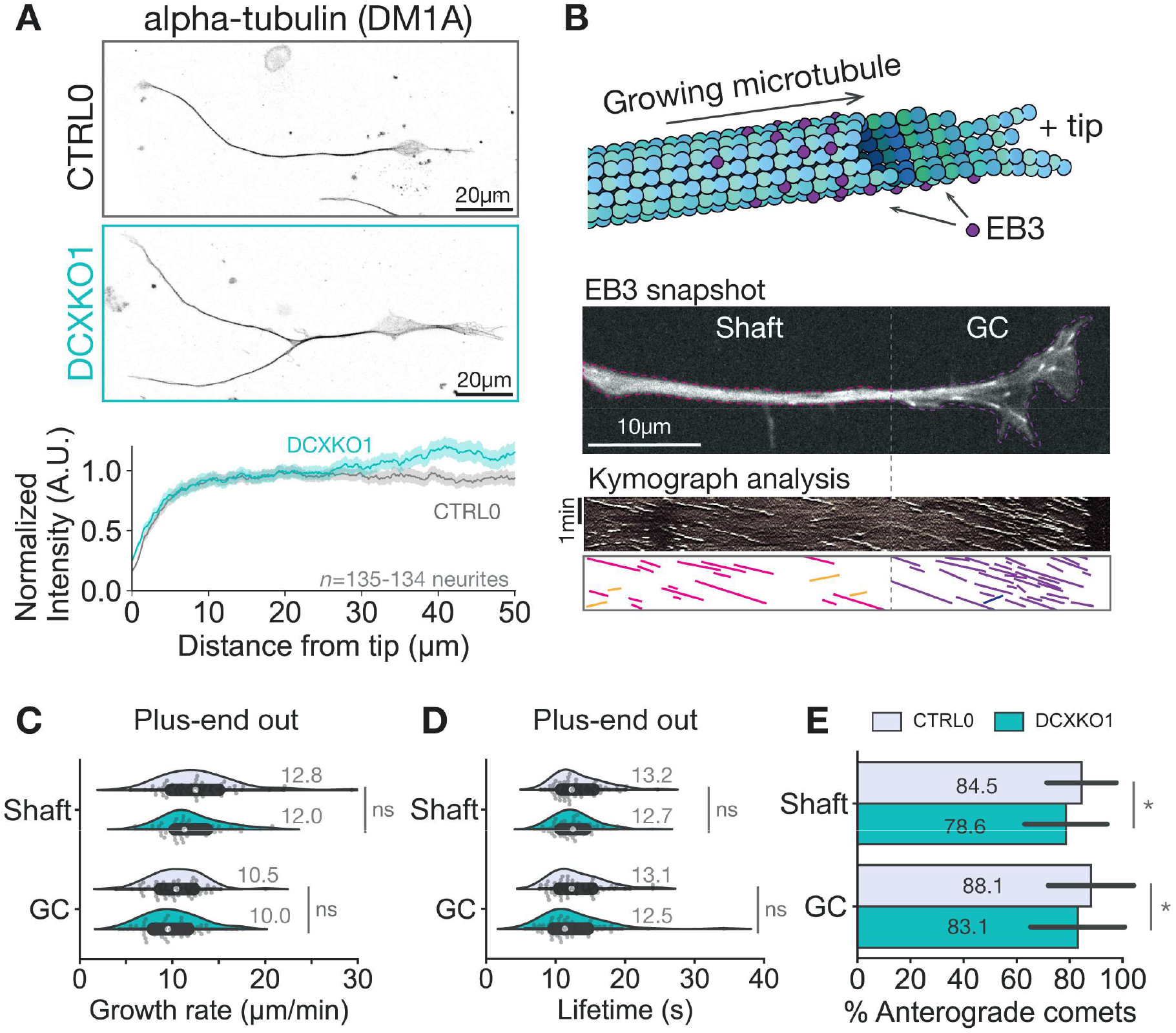
Microtubule dynamics are unaffected in DCX-KO neurons. **(A)** (Top) Immunostaining of CTRL0 and DCXKO1 neurons at DIV3 with a Tubulin-β3 specific antibody, and (Bottom) mean intensity profile found in the distal tip of neurons from both genotypes, SEM are shown in lighter grey (CTRL0) and lighter cyan (DCXKO1). Images included in the analysis are from *n*=135 CTRL0 neurites and *n*=134 DCXKO1 neurites from 3 experiments. Statistical analysis was performed using the Mann-Whitney non parametric U test: * *p*<0.05. **(B)** Schematics of the experimental procedure and analysis method for measuring microtubule dynamics. NPCs are electroporated with the plus-tip marker EB3-mCherry, and neurons are imaged at DIV3. Each comet is assigned to a domain (Shaft or growth cone (GC)), and its characteristics are measured: length, lifetime, growth rate, and direction. **(C)** Violin plot showing the distribution of growth rates for plus end out comets in shaft or growth cone of CTRL0 and DCXKO1 neurons. **(D)** Violin plot showing the distribution of lifetimes for plus end out comets in shaft or growth cone of CTRL0 and DCXKO1 neurons. **(E)** Box plot showing the proportion of plus end out comets in the shaft and growth cone of CTRL0 and DCXKO1 neurons. For C-E, *n*=63 CTRL0 and *n*=63 DCXKO cells from 4 experiments. Statistical analysis was performed using the Mann-Whitney non parametric U test: ns nonsignificant, * *p*<0.05.

We then looked specifically for changes in microtubule dynamics. We observed dynamic microtubules by transient expression of the end-binding protein EB3-mCherry in DCX-KO neurons (Fig. 3B and videos S7-S8). Since DCX is enriched in neurite tips (Fig. S4E), we measured neurite tips and shafts separately. However, regardless of region, we found no significant differences in EB3 comet dynamics between control and DCX-KO (Fig. 3C-D and Fig. S5), including in the densities of EB3 comets (0.40 ± 0.04 µm^-2^ control vs 0.44 ± 0.04 µm^-2^ DCXKO1, *p*=0.43), their velocities (10.5 ± 0.3 µm/min control vs 10.0 ± 0.4 µm/min DCXKO1, *p*=0.24), and lifetimes (GC: 13.1 ± 0.5 s control vs 12.5 ± 0.6 s DCXKO1, *p*=0.18). The characteristics of EB3 comet dynamics in our iPSC-derived neurons were equivalent to what has been reported in other studies (Lindhout et al. 2020; Dema et al. 2023). We did observe a slight increase in the proportion of retrograde comets, suggesting a minor role for DCX in the orientation of growing microtubules (Fig. 3E and Fig. S5E), consistent with recent cryo-electron tomography of microtubules in *Dcx*^*-/y*^ mouse neurons (Atherton et al. 2022). We also noticed that the area covered by EB3-decorated microtubules was reduced in DCX-KO neurites, indicating that DCX may contribute to growth cone morphology (Fig. S5C). Taken together, we conclude that DCX does not regulate EB3 comet dynamics in our model. Rather, other factors determine the growth rates, lifetimes, and nucleation rates in neurons. DCX-KO does appear to increase the rate of post-catastrophe shrinkage (Dema et al. 2024), which is consistent with reconstitution results(Moores et al. 2006). However, faster shrinkage rates in DCX-KO may not be sufficient to explain the increase in branch initiation events reported here (Fig. 2G).

### DCX-KO neurons show altered DCLK1 localization and lysosome motility

DCX has also been proposed to play a role in regulating intracellular trafficking, and dysfunctional cargo transport could be responsible for the observed excessive neurite branching. However, there is some ambiguity between observations made *in vitro* and in cells. For example, *in vitro*, DCX reduces the landing rate and run length of kinesin-1 motors but not kinesin-3 motors (Monroy et al. 2020), while in neurons, DCX shRNA caused reduced run lengths of kinesin-3 cargos (Liu et al. 2012), a result that may be confounded by off-target RNAi effects (Baek et al. 2014). In addition, DCX binds directly to dynein complexes and negatively regulates retrograde transport (Fu et al. 2022). The absence of DCX in our DCX-KO neurons might also modify the localization or behavior of other MAPs involved in the regulation of trafficking, such as Tau (Chaudhary et al. 2018; Monroy et al. 2018), MAP2 (Gumy et al. 2017), DCLK1 (Lipka et al. 2016), and others (Ramkumar, Jong, and Ori-McKenney 2018; Bodakuntla et al. 2019; Siahaan et al. 2022). As an example of this competition, DCX and Tau were shown to bind the microtubule lattice in a mutually exclusive manner in non-neuronal cell lines (Ettinger et al. 2016) and in rat neurons (Ettinger et al. 2016; Tint et al. 2009).

To determine how intracellular trafficking may be directly or indirectly impacted by DCX-KO, we measured the intensity profile of several molecular motors and MAPs in control and DCX-KO neurons by immunocytochemistry (Fig. 4A-B and Fig. S6). To maximize signal, we did not use detergent to extract the cytoplasmic pool of proteins (See methods, bound proteins only Fig. S6A) and detect both microtubule-bound and unbound proteins. The intensity of polarity markers MAP2 and Tau did not change in DCX-KO neurons, perhaps because they are not strongly expressed in neurons at this early pre-polarization stage (Lindhout et al. 2020). We found an increased signal by 1.4-fold for DCLK1 in DCX-KO neurons (Fig. 4A), which was not associated with higher expression levels (Fig. S6C). DCLK1 is enriched in the distal tip of neurites in our cell model (Fig. S6B), suggesting increased occupancy of the DCX binding site by its homologue (Tanaka, Koizumi, and Gleeson 2006; Deuel et al. 2006). Kinesin-1, kinesin-2, kinesin-3 and dynein intensities were not significantly modified in DCX-KO neurons (Fig. 4A and Fig. S6A). However, the lack of change in motor localization (Fig. S6B) does not exclude a defect in cargo trafficking.

**Figure 4:**
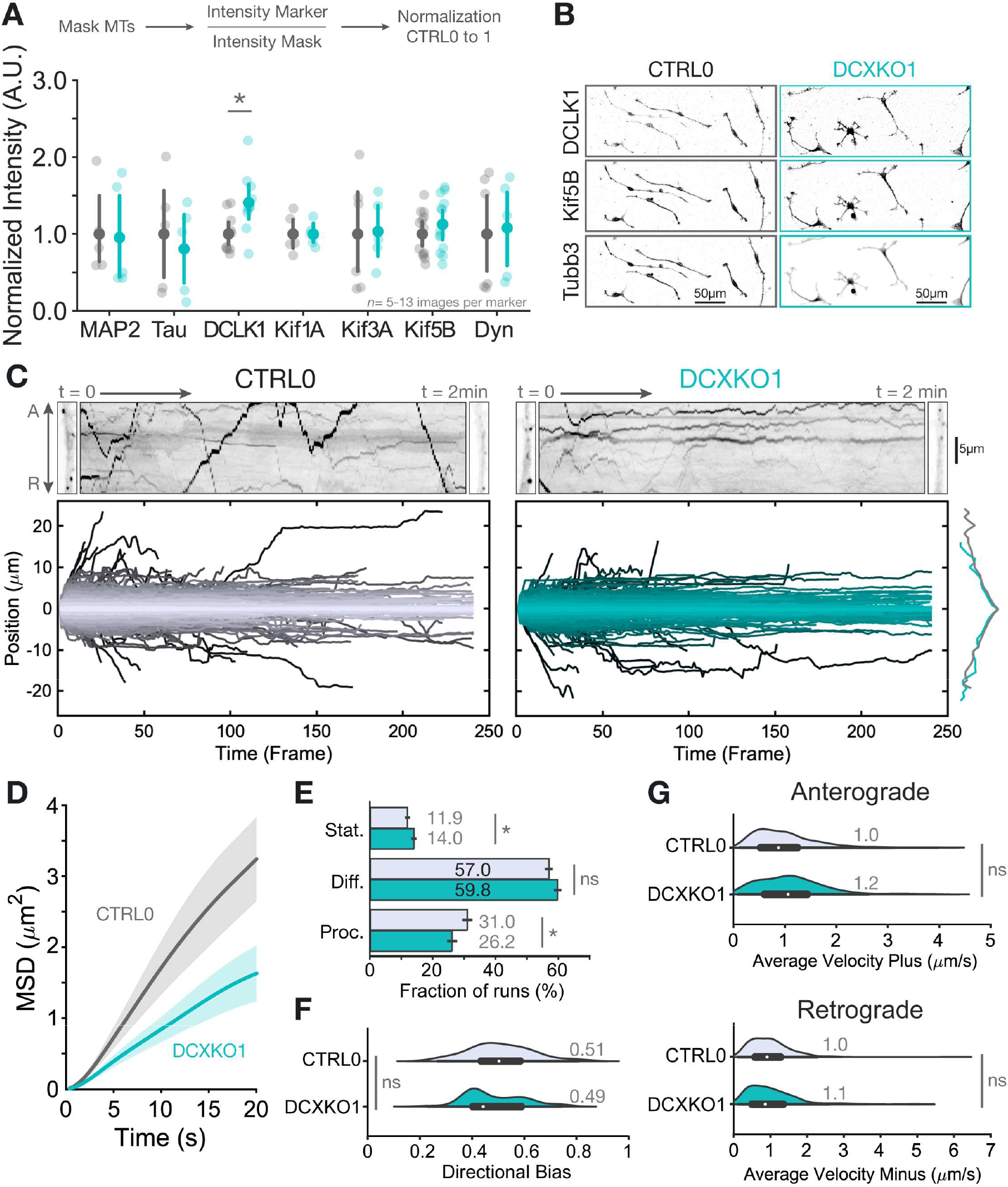
DCX-KO neurons show altered DCLK1 localization and lysosome motility. **(A)** Compared mean intensity for MAP2, Tau, DCLK1, Kif1A, Kif3A, Kif5B, and Dynein in CTRL0 or DCXKO1 DIV3 immuno-stained neurons. The analysis pipeline is shown above the plot: a mask of the microtubules (MTs) was made, and the intensity of the marker was normalized to the intensity of the mask and to the control condition. Each transparent point represents a single image, while the opaque points show mean values, error bars represent the confidence intervals. Analyses are from *n*=5-13 CTRL0 or DCXKO1 images from 2-4 experiments. Statistical analysis was performed using the Mann-Whitney non parametric U test: * *p*<0.05. **(B)** Representative images from CTRL0 and DCXKO1 DIV3 neurons stained for DCLK1, Kif5B and Tubb3. **(C)** (Top) Representative still images of CTRL0 and DCXKO1 straight neurite sections after incubation with the Lysotracker® dye, and 2min kymographs of those same sections. A means anterograde, and R retrograde. (Bottom) Position plots for lysosomes detected using Trackmate during the 241 frames for CTRL0 and DCXKO1 neurons. The distribution of these positions was also plot and is represented on the right end side of the position plots. **(D)** Mean squared displacement (MSD) of lysosomes inside CTRL0 and DCXKO1 neurites. **(E)** Proportion of tracked cargos that are either stationary (Stat.), Diffusive (Diff.) or Processive (Proc.). **(F)** Directional bias of lysosome cargos in CTRL0 and DCXKO1 neurons, where 0 corresponds to retrograde only and 1 corresponds to anterograde only. **(G)** Violin plots of the average velocity for anterograde (plus-ended) displacement and retrograde (minus-ended) displacement of lysotracker® cargos in CTRL0 and DCXKO1 DIV3 neurons. All trajectories (6082 for control and 3887 for DCXKO1) were obtained from *n*=41 control and 30 DCXKO1 neurons from 3 independent experiments. Statistical analysis was performed using the Student’s t-test: ns nonsignificant, * *p*<0.05.

To test the role of DCX in regulating cargo trafficking, we measured the trajectories of lysosomes using the pH-sensitive Lysotracker® dye (Videos S9-S10). Lysosomes move in a bidirectional manner using kinesin-1, kinesin-2, kinesin-3 and dynein motors, making them well-suited to assess broad trends in intracellular trafficking (Pu et al. 2016; Bentley et al. 2015). Figure 4C shows Lysotracker kymographs and extracted trajectories for control and DCX-KO neurons. In both cases, we observed bidirectional transport with similar distributions of trajectories (right panel). We characterized this transport further by measuring the mean-squared displacement (MSD) of lysosome position, radius of gyration (Rg), directional bias, and velocities as a function of time (Fig. 4D-G and S7). The Rg, MSD and extracted α value (See methods: slope of the MSD plot) which assess general motility of cargos are significantly decreased in DCX-KO neurons (Fig. 4D and Fig. S7A-B), confirming the impact of the KO on intracellular trafficking. All the trajectories used for MSD assessment can be sorted as stationary, diffusive or processive depending on their characteristics (Fig. 4E, see Methods). Interestingly, the proportion of processive motility is significantly reduced in DCX-KO neurons (control 31.0 ± 1.3 % vs DCX-KO 26.2 ± 1.3 %, *p*=0.012, *) and correspondingly the fraction of diffusive (control 57.0 ± 1.0 % vs DCX-KO 59,8 ± 0.9 %, *p*=0.053, ns) and stationary motility increased (control 11,9 ± 0.5 % vs DCX-KO 14,0 ± 0.6 %, *p*=0.010, *).

We noticed that lysosomes in DCX-KO neurons are more unidirectional than in control cells, as indicated by a bimodal directional bias (Fig. 4F). However, lysosome velocity was not affected in either the minus- or plus-end direction (Fig. 4G). Altogether, in DCX-KO neurons, lysosomes are less processive. These results support the principle that DCX-KO impacts intracellular trafficking, and a comprehensive analysis of neuronal cargos will be an important future direction of research. However, the defect in lysosome trafficking is mild and possibly insufficient to explain the excessive branching phenotype found in DCX-KO neurons (Fig. 2).

Thus far, our analysis of DCX-KO neurons has revealed complex and pleiotropic phenotypes, namely a slowed nucleokinesis, increased rate of branch initiation, increased proportion of misoriented microtubules, and reduction in lysosome processivity. These disparate phenotypes point to an upstream regulator of microtubule physiology: something which, if dysregulated, disrupts microtubule homeostasis at many levels. We hypothesized that this upstream regulator could be the tubulin code.

### DCX-KO disrupts microtubule polyglutamylation

The tubulin code is an information system for microtubule specialization that contributes to the processivity of motor proteins (Sirajuddin, Rice, and Vale 2014; Tas et al. 2017), binding affinity of MAPs (Boucher et al. 1994; Bonnet et al. 2001), and microtubule spacing and geometry (Cueva et al. 2012). For example, acetylated microtubules (Acetyl) are more flexible and predominantly bundled (Portran et al. 2017; Balabanian, Berger, and Hendricks 2017; Tas et al. 2017). Recently, acetylation has been linked to neurite branching in a model of an α-tubulinopathy (Hoff et al. 2022). Tyrosinated microtubules (Tyr) act as preferred tracks for kinesin-3, kinesin-5 (Sirajuddin, Rice, and Vale 2014; Lessard et al. 2019), and dynein (McKenney et al. 2016; Tas et al. 2017); Polyglutamylation (polyE) and detyrosination (detyr) regulate MCAK and severing enzymes binding, that are responsible for modified branching patterns when depleted (Homma et al. 2003; Valenstein and Roll-Mecak 2016), reviewed in (Moutin et al. 2021; Janke and Magiera 2020). Our results pointed to many of these proteins and pathways. Therefore, we wondered if the tubulin code could be altered in DCX-KO neurons.

To test for changes in the tubulin code, we performed immunocytochemistry and Western blots in control and DCX-KO neurons, with antibodies against acetylation (611B1), tyrosination (YL1/2), detyrosination (detyr), initial glutamylation (GT335) and polyglutamylation (polyE) where glutamate side chains are reversibly added to the C-terminal tails of α- and β-tubulin (Audebert et al. 1993; Redeker et al. 1992). (Fig. 5A-B and S8). GT335 recognizes all glutamate chains through the initial residue, while polyE detects glutamate chains of 4 residues or more (Wolff et al. 1992; Lacroix et al. 2010). Remarkably, we observed a dramatic reduction specifically in tubulin polyglutamylation by both Western blot and immunocytochemistry (polyE – Fig. 5A-B and Fig. S8). In addition, detection of all glutamate chains with GT335 in Western blots is mildly increased in DCX-KO neurons. We analyzed the subcellular distribution of staining in neurites and cell bodies (CB) and confirmed an increase in initial glutamylation specific to cell bodies and a drastic reduction in polyglutamylation only in neurites (Fig. 5C-D). In parallel, we observed that acetylation and detyrosination levels, which label long-lived microtubules, were unchanged in DCX-KO neurons compared to control, while tyrosination is mildly increased in cell bodies. These results indicate that DCX expression regulates the tubulin code, especially polyglutamylation levels in neurites. The reduced levels of polyglutamylation in DCX-KO will impact a broad spectrum of MAPs and motors.

**Figure 5:**
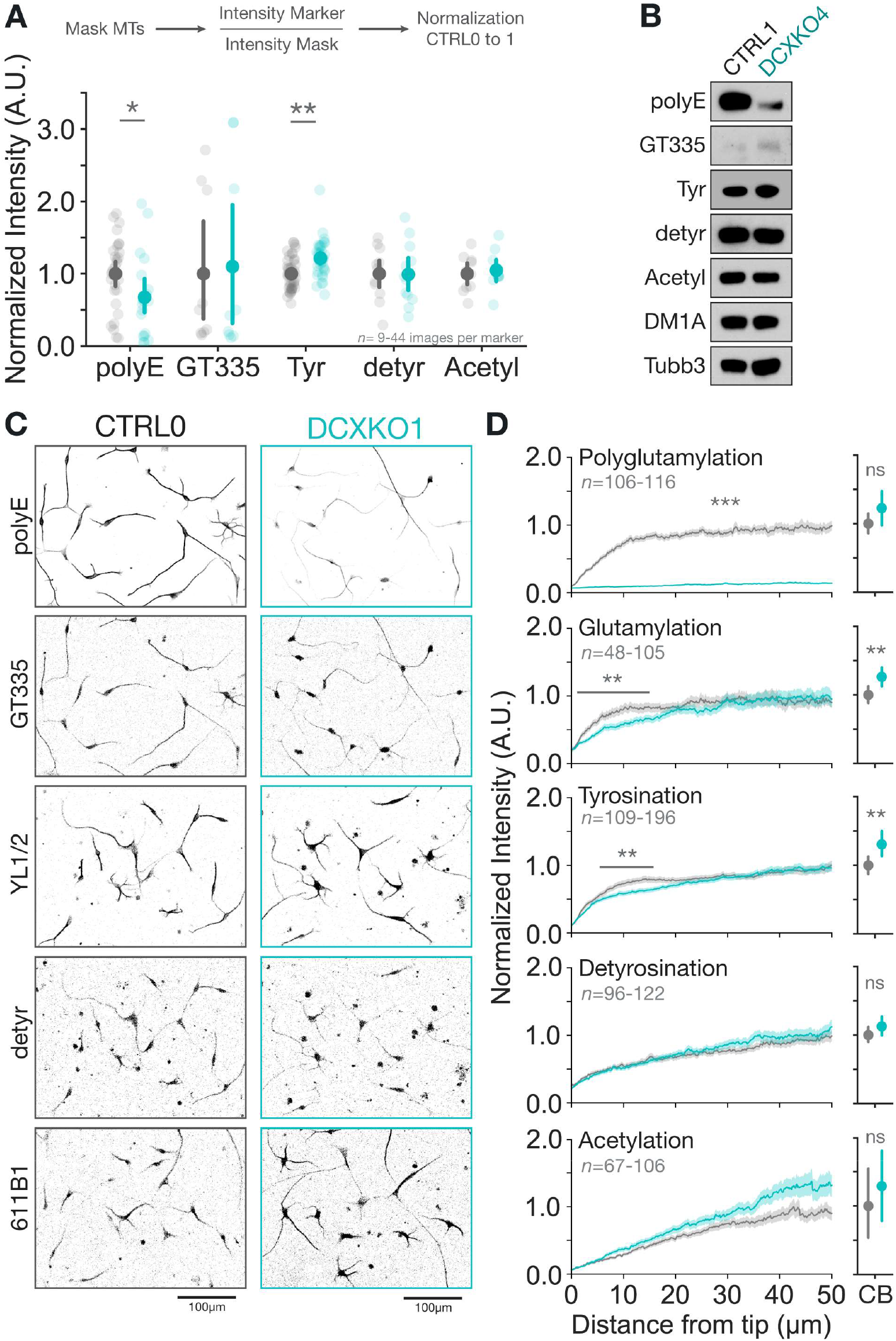
DCX-KO disrupts microtubule polyglutamylation. **(A)** Compared mean intensity for polyglutamylation (polyE), glutamylation (GT335), tyrosination (Tyr), detyrosination (detyr), and acetylation (Acetyl) in CTRL0 or DCXKO1 DIV3 immunostained neurons. The analysis pipeline is shown above the plot: a mask of the microtubules (MTs) was made, and the intensity of the marker was normalized to the intensity of the mask and to the control condition. Each transparent point represents a single image, while the opaque points show mean values, error bars represent the confidence intervals. Plots are from *n*=9-44 CTRL0 or DCXKO1 images from 3-6 experiments. Statistical analysis was performed using the Mann-Whitney non parametric U test: * *p*<0.05, ** *p*<0.01. **(B)** Immunoblots showing the relative post-translational modifications levels in CTRL1 and DCXKO4 DIV3 neurons. The same samples were loaded on different gels for each antibody. **(C)** Representative images of CTRL0 and DCXKO1 neurons stained for PTMs with the following antibodies: polyE, GT335, YL1/2, detyr, 611B1. **(D)** Intensity profiles for the PTMs levels in distal neurites and cell bodies (CB) of CTRL0 and DCXKO1 DIV3 neurons. Error bars represent the confidence interval. Plots are from *n*=48-196 CTRL0 or DCXKO1 neurites and *n*=23-90 cell bodies, from at least 3 experiments. Statistical analysis was performed using the Mann-Whitney non parametric U test: ns nonsignificant, * *p*<0.05, ** *p*<0.01, *** *p*<0.001.

### DCX reduces neurite density by increasing tubulin polyglutamylation levels

We next asked if the regulation of PTM levels by DCX could result in the branching phenotype we have seen in DCX-KO neurons. To mechanistically link DCX and polyglutamylation, we tested whether the expression of DCX could rescue polyglutamylation levels and the branching phenotype in DCX-KO neurons. We electroporated a DCX-GFP construct in control and DCX-KO neurons and measured both the polyglutamylation (polyE) levels and neurite density (Fig. 6A-C and Fig. S9A). In DCX-KO neurons re-expressing DCX-GFP, we found that polyglutamylation levels were increased and neurite density was accordingly reduced to match control neurons. Surprisingly, the overexpression of DCX-GFP in control neurons reduced polyglutamylation levels and increased neurite density, similar to DCX-KO neurons (Fig. 6B-C). This result is consisent with overexpression experiments done in mouse models, which also led to an increase in neurite numbers (Cohen, Segal, and Reiner 2008; Yap et al. 2016). Moreover, our data indicates that the effects of DCX protein concentrations are biphasic, where an intermediate protein concentration achieves healthy polyglutamylation levels and neurite densities, while reduced or elevated levels of DCX lead to dysregulation.

**Figure 6:**
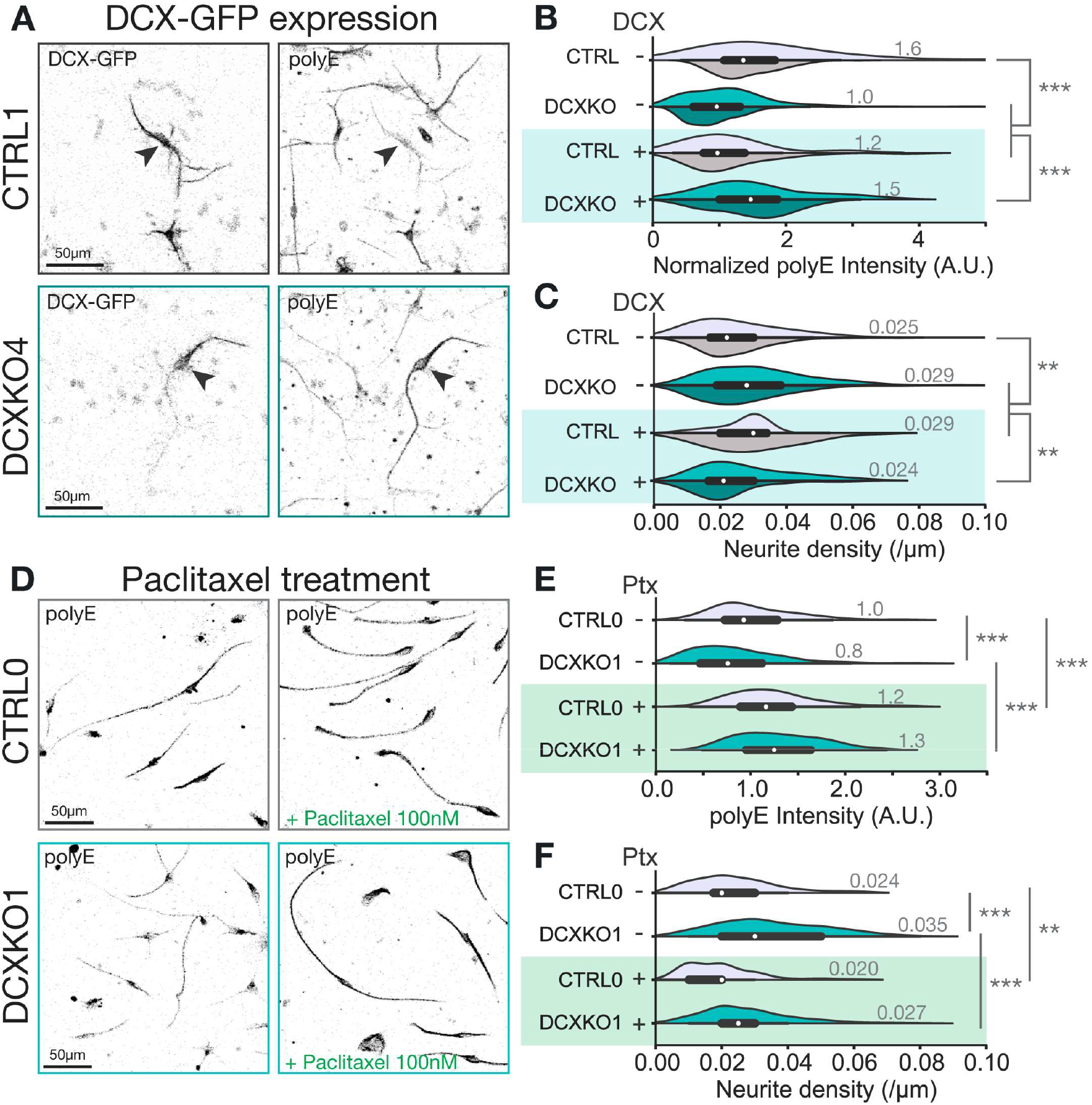
DCX reduces neurite numbers by increasing tubulin polyglutamylation levels. **(A)** Representative images of CTRL1 and DCXKO4 DIV3 neurons electroporated with DCX-GFP and stained for polyglutamylation. The arrowheads are showing the same cell in both channels (GFP, and polyE). **(B)** Violin plots showing the polyglutamylation (polyE) intensity normalized by Tubb3 intensity in CTRL and DCXKO DIV3 neurons with or without DCX-GFP electroporation. **(C)** Violin plot showing the neurite density in CTRL and DCXKO DIV3 neurons with or without DCX-GFP electroporation. For B-C, the analysis included *n*=97 CTRL0 (light grey), 178 CTRL1 (dark grey), 225 DCXKO1 (cyan), 171 DCXKO4 (dark cyan), 11 CTRL0+DCX-GFP, 34 CTRL1+DCX-GFP, 38 DCXKO1+DCX-GFP, and 45 DCXKO4+DCX-GFP neurons. Each line was used in 3 paired experiments. These two plots were cut-off to better show the main population, but the maximum values for polyE intensity, and neurite density are 9.4 (A.U.), 0.156 neurite/µm respectively. Statistical analysis was performed using the Mann-Whitney non parametric U test: ns nonsignificant, ** p<0.01, *** p<0.001. **(D)** Representative images of CTRL0 and DCXKO1 DIV3 neurons stained for polyglutamylation (polyE) after 48h in regular media or 48h in media supplemented with 100nM taxol. **(E)** Violin plots showing the polyglutamylation (polyE) intensity in CTRL0 and DCXKO1 DIV3 neurons with or without treatment with taxol. **(F)** Violin plots showing the neurite density in CTRL0 and DCXKO1 DIV3 neurons with or without treatment with taxol. For E-F, images included in the analysis are from *n*=84 CTRL0, 172 DCXKO1, 120 CTRL0+taxol and 160 DCXKO1+taxol neurons from 2 experiments. Statistical analysis was performed using the Mann-Whitney non parametric U test: ns nonsignificant, ** *p*<0.01, *** *p*<0.001.

### Chemical induction of polyglutamylation by paclitaxel restricts neurite density

As a first attempt to rescue polyglutamylation levels and neurite density, we treated cells with paclitaxel, a well-known chemotherapy drug, that stabilizes microtubules (Yang and Horwitz 2017). Low doses of paclitaxel were previously shown to increase PTM levels by promoting detyrosination, acetylation, and polyglutamylation (Hammond et al. 2010). We therefore applied paclitaxel (100 nM) for 48h to neurons and measured their polyglutamylation levels and neurite density (Fig. 6D-F and Fig. S9B-D). Neurons treated with paclitaxel showed an increase in polyglutamylation levels in their branches independent of their genotype (Fig. 6E). Polyglutamylation was increased by ∼1.2-fold in control cells and by 1.6-fold in DCX-KO cells. In parallel to these changes in polyglutamylation levels, neurite density decreased by about 20% in paclitaxel-treated cells (Fig. 6F), indicating a correlation between polyglutamylation levels and neurite density. However, because paclitaxel affects multiple PTMs and causes lattice expansion (Arnal and Wade 1995; Alushin et al. 2014), we sought a more targeted approach to rescue polyglutamylation in our DCX-KO system.

### Expression of the α-tubulin elongase TTLL11 in DCX-KO neurons rescues neurite density

Polyglutamylation levels are determined by modifying enzymes, and we hypothesized that the transient expression of a glutamylating enzyme could also rescue polyglutamylation levels together with neurite density in our model. A family of 9 glutamylases (Tubulin Tyrosine Ligase Like, or TTLLs) (Janke et al. 2005; van Dijk et al. 2007) competes against a family of 6 deglutamylases (Cytosolic CarboxyPeptidase, or CCPs) (Rogowski et al. 2010) to create glutamate chains of variable length and structure (Janke and Magiera 2020). The members of each family have specialized roles, such as branch initiation (e.g., TTLL4, TTLL7) versus elongation (e.g., TTLL6, TTLL11) (Mahalingan et al. 2020), or shortening (e.g., CCP1, CCP6) versus branch removal (CCP5) (Rogowski et al. 2010). Additionally, α-tubulin and β-tubulin polyglutamylation are distinct (Edde et al. 1990; Alexander et al. 1991; Bodakuntla et al. 2021), with elongase TTLL’s showing preference for α-tubulin. We confirmed that our neurons show exclusively α-tubulin polyglutamylation (Fig. S10A), consistent with previous studies of early differentiated neurons (Magiera and Janke 2013). Thus, we choose to express an α-tubulin specific elongase, namely TTLL11. As one of the main autonomous enzyme expressed in neurons (McKenna et al. 2023), TTLL11 is able to elongate pre-existing α-tubulin glutamate chains (van Dijk et al. 2007).

Because TTLL11-mRuby expression in neurons was barely measurable after fixation and staining (data not shown), we modified our electroporation protocol to include a sorting step (See methods and Fig. S10B). Figure 7A shows representative images of polyglutamylation levels in control and DCX-KO neurons with (mRuby +) or without (mRuby -) TTLL11-mRuby expression. In both control and DCX-KO neurons expressing TTLL11-mRuby, polyglutamylation levels increased about 5-fold (Fig. 7B and Fig. S10E-F). In parallel, we found that the expression of TTLL11 in DCX-KO neurons rescued neurite density back to control levels (Fig. 7C and Fig. S10C-D). These results match DCX expression and paclitaxel treatments in neurons (Fig. 6), where DCX is sufficient but not required to control polyglutamylation levels. We can therefore confirm that DCX restricts neurite branching through regulation of polyglutamylation levels.

**Figure 7:**
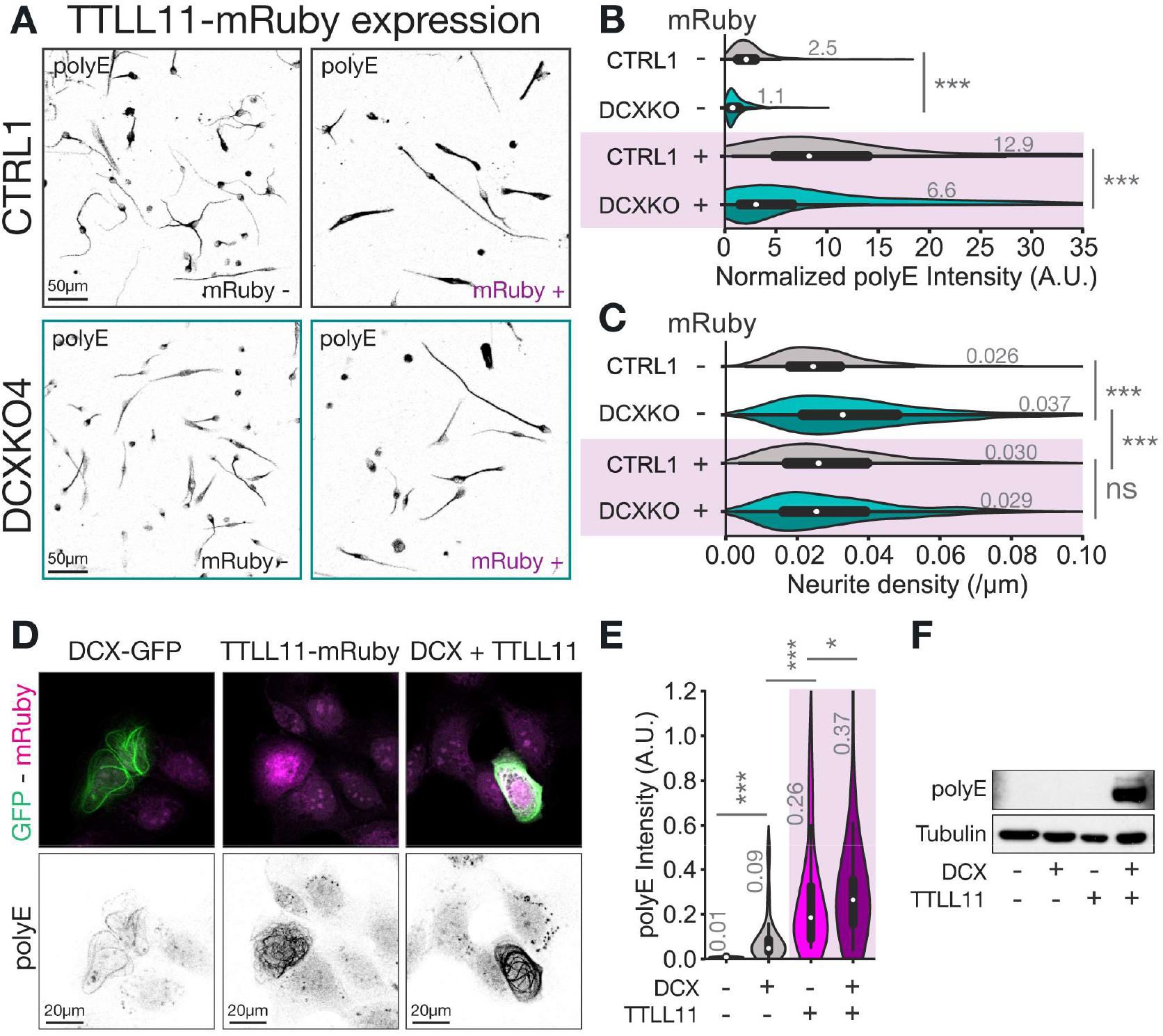
DCX and TTLL11 synergize to regulate polyglutamylation levels. **(A)** Representative images of CTRL1 and DCXKO4 DIV3 neurons control (mRuby-) or electroporated with TTLL11-mRuby (mRuby +) and stained for polyglutamylation (polyE). **(B)** Violin plots showing the polyglutamylation (polyE) intensity in CTRL1 and DCXKO DIV3 neurons with (+ or purple area) or without (-) TTLL11-mRuby electroporation. DCXKO1 and DCXKO4 are shown as split violins, with DCXKO1 in cyan (top) and DCXKO4 in dark cyan (bottom). **(C)** Violin plot showing the neurite density in CTRL1 and DCXKO DIV3 neurons with or without TTLL11-mRuby electroporation. For B-C, images included in the analysis are from n=372 CTRL1 (dark grey), 380 DCXKO1 (cyan), 654 DCXKO4 (dark cyan), 199 CTRL1+TTLL11-mRuby, 186 DCXKO1+TTLL11-mRuby and 408 DCXKO4+TTLL11-mRuby neurons from 6 independent experiments. These two plots were cut-off to better show the main population, but the maximum values for polyE intensity, and neurite density are 91.6 (A.U.), 0.143 neurite/µm respectively. Statistical analysis was performed using the Mann-Whitney non parametric U test: ns nonsignificant, *** p<0.001. **(D)** Representative images of U2OS cells transfected respectively with DCX-GFP (green), TTLL11-mRuby (magenta) or DCX-GFP and TTLL11-mRuby together. All conditions were fixed and stained for polyglutamylation (polyE). **(E)** Violin plots showing the quantification of polyglutamylation (polyE) levels in U2OS transfected cells as in D. Images included in the analysis are from n=27 control (untransfected), 58 DCX-GFP, 46 TTLL11-mRuby and 95 DCX+TTLL11 cells from 3 experiments. This plot was cut-off to better show the main population, but the maximum values for polyE intensity is 2.46 (A.U.). Statistical analysis was performed using the Mann-Whitney non parametric U test: * *p*<0.05, *** *p*<0.001. **(F)** Representative immunoblot of U2OS cells untransfected or transfected respectively with DCX-GFP, TTLL11-mRuby or DCX+TTLL11, and stained for polyglutamylation (polyE) and tubulin (DM1A). DCX and TTLL11 transfections were visually checked before lysis of the samples. The same samples were loaded on multiple gels for each antibody.

### DCX and TTLL11 synergize to regulate polyglutamylation levels

The rescue of DCX-KO phenotypes by TTLL11 expression caused us to wonder if DCX is able to regulate polyglutamylation by direct activation of TTLL11. To better understand the relationship between these two proteins, we used U2OS cells as a simplified model system. As with other cancer lines, U2OS cells have low TTLL expression and thus minimal polyglutamylation levels at interphase (Wolff et al. 1992; van Dijk et al. 2007; Zadra et al. 2022). In addition, U2OS cells do not express DCX, which is specific to developing neurons. The use of this model allowed us to isolate the effect of DCX and TTLL11 expression on polyglutamylation levels. As illustrated in figure 7D, U2OS cells were transfected either with DCX-GFP (green), TTLL11-mRuby (magenta) or both constructs together, and their polyglutamylation levels were measured by immunocytochemistry and Western blots (Fig. 7D-F and Fig. S11). Both methods revealed equivalent results: untransfected cells had no detectable polyE signal, while cells expressing DCX, TTLL11 or both showed different degrees of polyglutamylation (Fig. 7E-F and Fig. S11C), with signal clearly localizing to microtubules (Fig. 7D). DCX expressing cells had 12 times more polyglutamylation than control: DCX is not a glutamylase, so its expression alone likely affects the modifying enzymes naturally present in the cells to trigger polyglutamylation. TTLL11 expressing cells had 36 times more polyglutamylation than control: the expression of TTLL11 alone is sufficient to shift the balance of modifying enzymes, as in our neuronal model. In coexpressing cells, only lower levels of DCX and TTLL11 expression were achieved (35% less DCX and 30% less TTLL11, Fig. S11A). However, the polyglutamylation produced by co-expression was 53 times more than control (Fig. 7E). Thus, polyglutamylation levels are 60% higher than expected from the addition of DCX alone and TTLL11 alone activity (Fig. S11B). These results show that DCX and TTLL11 work synergistically to control polyglutamylation in U2OS cells. We found similar results in neurons, as polyglutamylation levels were 40% higher than expected from the addition of DCX-induced and TTLL11-induced glutamylase activity (Fig. 7B). Altogether, these experiments state DCX as a regulator of neurite density through control of tubulin polyglutamylation, and more specifically activation of glutamylases like TTLL11.

### DCX patient-mutant neurons have reduced polyglutamylation levels

To test the relevance of our findings in the context of disease, we used our CRISPR/Cas9 approach to introduce lissencephaly-causing patient mutations into DCX in iPSCs. We used previously characterized missense mutations located in both domains of DCX: S47R, Y64N, D86H in DC1 and R178L, P191R, and R192W in DC2 (Bahi-Buisson et al. 2013; Leger et al. 2008; Bechstedt, Lu, and Brouhard 2014). We assessed polyglutamylation levels in neurons derived from these isogenic mutated iPSC lines by Western blot and immunocytochemistry (Fig. 8A-B). Remarkably, all patient mutations tested resulted in reduced polyglutamylation levels independent of the mutation locus. Overall, these findings demonstrate a central role for DCX in regulating polyglutamylation levels and thus MAP affinity and microtubule organization.

**Figure 8:**
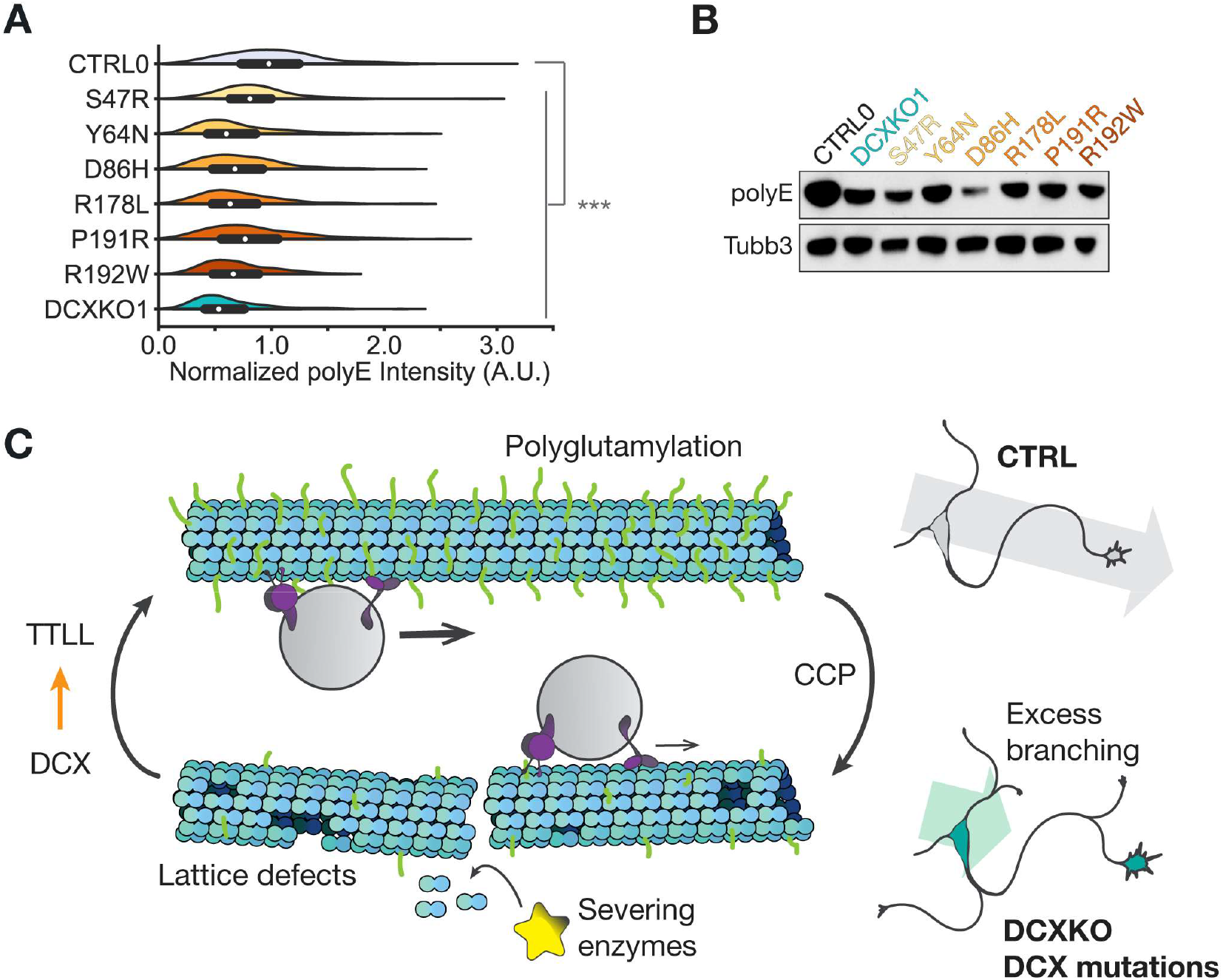
Model of a feedback regulation of the tubulin code by the MAP DCX. **(A)** Violin plots showing the polyglutamylation (polyE) intensity in stained DIV3 neurons carrying the following DCX patient mutations: S47R, Y64N, D86H, R178L, P191R, R192W, and compared to CTRL0 and DCXKO1. Images included in the analysis are from *n*= 384 CTRL0, 314 D86H, 177 DCXKO1, 323 P191R, 260 R178L, 287 R192W, 215 S47R, 289 Y64N neurons from 3 experiments. Statistical analysis was performed using the Mann-Whitney non-parametric U test: *** *p*<0.001. **(B)** Immunoblots showing polyglutamylation levels in CTRL0, DCXKO1 and mutant DIV3 neurons (DC1: S47R, Y64N, D86H, DC2: R178L, P191R, R192W). The same samples were loaded on different gels for both antibodies. **(C)** Control neurons have high polyglutamylation levels. In this context, DCX activates TTLLs leading to two molecular phenotypes in DCX-KO neurons: a decrease in polyglutamylation of the microtubule lattice and an impeded cargo motility. The addition or association of these two mechanisms result in an excessive production of neurite and a decrease in migration speed and efficacy in DCX-KO and mutated neurons.

## Discussion

In summary, using novel DCX knockout iPSC model, an orthogonal U2OS system, and iPSC models with disease mutations, we show that DCX regulates α-tubulin polyglutamylation to restrict neurite branching. We believe these results provide insight into the cellular basis of double-cortex syndrome and X-linked lissencephaly. DCX-KO neurons showed pleiotropic defects, including meandering nucleokinesis (Fig. 1), excessive branch initiation (Fig. 2), and impaired lysosome trafficking (Fig. 4), but without changes in EB3 comet dynamics (Fig. 3). Rather, our results indicated that these pleiotropic defects originate from a strong reduction in α-tubulin polyglutamylation levels (Fig. 5), an unexpected finding not predicted in the literature on DCX (Fig. 8). Both polyglutamylation levels and neurite density were rescued by DCX expression, by paclitaxel treatment (Fig. 6), and by TTLL11 expression (Fig. 7), indicating a robust connection between DCX and the tubulin code. These results contribute to a growing body of evidence on the importance of polyglutamylation for morphology in many types of neurons (Gavoci et al. 2024; Lopes et al. 2020; Ziak et al. 2024).

How could DCX help maintain a healthy level of polyglutamylation in developing neurites? Polyglutamylation is a complex, reversible, and heterogenous feature of neuronal microtubules. The patterns of polyglutamylation are generated by the competition between glutamylases (TTLLs) and deglutamylases (CCPs), proteins whose regulation is essential for neuronal health. Excessive polyglutamylation upon loss of CCP activity causes neurodegeneration (Magiera et al. 2018; Shashi et al. 2018), while our results link reduced polyglutamylation to a neuronal migration disease. DCX could contribute to a healthy level between these extremes by stimulating the activity of TTLL’s (more specifically, α-tubulin elongases) or by reducing the activity of CCP’s. At present, our data provides evidence for a model based on TTLL activation, as DCX and TTLL11 co-expression caused a synergistic increase in polyglutamylation levels in U2OS cells (Fig. 7D-F). However, it is important to note that we cannot yet rule out inhibition of CCP’s, which could occur concurrently. Because mono-glutamylation is preserved in our model (Fig. 4D), the relevant CCP’s would be chain-shortening enzymes (CCP1, CCP2, CCP3, CCP4, CCP6), not de-branching enzymes (CCP5). In either case, regulation of long α-tubulin polyglutamate chains (as detected by polyE antibodies) explains how DCX restricts the number of neurite branches (Fig. 8C). Interestingly, it has been shown previously that TTLLs are present at low levels at this stage of maturation (Lindhout et al. 2020) have low microtubule affinity (van Dijk et al. 2007), and are enriched mainly in the cell bodies of neurons (O’Hagan et al. 2017) rather than the neurites.

Thus, neurites appear to establish healthy polyglutamylation levels using a relatively small number of polyglutamylases. In this context, DCX may play a key role as a positive activator of TTLL-microtubule interactions.

We can imagine two mechanisms by which DCX could regulate TTLLs and cause, e.g., the synergistic effect observed in U2OS cells. First, DCX could directly bind the TTLL enzymes to bring them to the microtubule lattice, e.g. by direct recruitment of the conserved cationic microtubule-binding domain of TTLLs (Garnham et al. 2015). The cationic domain of the TTLL6 bound to the α-tubulin tail shows a helix extending into the interprotofilament groove (Mahalingan et al. 2024), which the authors note is near the binding site of DCX, EB proteins, as well as kinetochore components. Second, DCX could indirectly affect the TTLL enzymes by changing the structural feature(s) of αβ-tubulins in the lattice, in other words by altering the enzymes’ substrate in a way that accelerates the enzyme’s kinetics. Along these lines, recent work has linked other PTMs to the structural feature of “lattice spacing”, or the measured dimer rise of the microtubule lattice, which varies between 81-84 Å in cryo-electron microscopy (LaFrance et al. 2022; Zhang, LaFrance, and Nogales 2018). More specifically, the detyrosination complex VASH/SVBP is more active on expanded, GTP-like microtubules lattice (Yue et al. 2023), as is the acetyltransferase αTAT1 (Shen and Ori-McKenney 2024). A cognate model based on “altered substrates” makes sense for DCX, given DCX’s sensitivity to structural features such as lattice curvature (Bechstedt, Lu, and Brouhard 2014), protofilament number (Moores et al. 2004; Bechstedt and Brouhard 2012), and lattice spacing (Manka and Moores 2018). However, an altered substrate model is not incompatible with direct recruitment, and detailed biophysical analysis will be needed to define these models and distinguish between them.

Our observation that DCX regulates polyglutamylation to restrict neurite branching (Fig. 6-7) harmonizes with many recently-discovered links between MAPs involved in branching pathways and the tubulin code. Consider the severing enzymes, spastin and katanin (Costa and Sousa 2022). The severing rate of these enzymes is increased by long polyglutamate chains (Lacroix et al. 2010) and sensitive to both α-tail and β-tail modifications (Genova et al. 2023). Reconstitution experiments indicated that the severing rate first increases as glutamate chains get longer but then decreases beyond some optimal chain length *in vitro* (Lacroix et al. 2010; Valenstein and Roll-Mecak 2016). Severing enzymes were originally proposed to fragment the microtubule network at branch points (Roll-Mecak and Vale 2006), such that the proliferation of short microtubules would stabilize transient actin protrusions (Yu et al. 2008). But fragmentation is not the only outcome of severing enzyme activity in the presence of soluble tubulin (Vemu et al. 2018); rather, severing enzymes create holes and defects in the lattice that are subsequently repaired by soluble GTP-tubulin. This process of microtubule “self-repair” (Schaedel et al. 2015) may refresh the lattice with respect to nucleotide state, PTMs, etc. Whether via fragmentation or damage, severing enzymes will be impacted by the reduction in polyglutamylation in DCX-KO neurons. A link between microtubule damage and DCX is hinted at by cryo-electron tomography of microtubules in Dcx^-/y^ mouse neurons, in which the number of large lattice defects increased relative to control (Atherton et al. 2022). To further complicate the picture, severing enzymes are inhibited by structural MAPs such as tau (Tan et al. 2019; Siahaan et al. 2022), and tau’s envelope formation on microtubules (Tan et al. 2019) is increased by polyglutamylation (Genova et al. 2023). The dynamics of this complex multi-protein system will be exciting to measure and to model.

Healthy polyglutamylation levels are important for motor proteins (Hammond et al. 2010; Sirajuddin, Rice, and Vale 2014), but DCX also regulates motor proteins directly, potentially having complex effects. In reconstitution assays, kinesin-1 has a reduced run length on polyglutamylated microtubules (Genova et al. 2023), and DCX-coated microtubules reduce kinesin-1’s landing rate (Monroy et al. 2020). Similarly, the run length of kinesin-3 is increased on polyglutamylated microtubules (Lessard et al. 2019), while DCX-coated microtubules do not affect kinesin-3 motility *in vitro* (Monroy et al. 2020). Both kinesin-1 and kinesin-3 are present on lysosomes (Guardia et al. 2016), and our results indicate that DCX-KO primarly affects the fraction of lysosomes which are stationary versus processive. Other cargos regulated by DCX-family proteins are synaptophysin (Deuel et al. 2006), Vamp2 (Liu et al. 2012), dense-core vesicles (Lipka et al. 2016), and TrkB (Fu et al. 2022). The phenotypes of disrupted intracellular trafficking will need to be explained in light of both direct interactions between DCX and motor proteins (Fu et al. 2022; Liu et al. 2012) as well as changes in polyglutamylation, and polyglutamylation’s effects on affinity of the other MAPs like tau (Genova et al. 2023) (Fig. 8).

Our study did not address a potential role for DCX in the regulation of the actin cytoskeleton. The data on DCX and actin are mixed. DCX has been proposed to modulate actin through interactions with spinophilin, an actin-binding protein (Bielas et al. 2007). Dcx^-/y^ neurons showed reduced actin staining in some experiments (Fu et al. 2013), but this result was not reproduced in recent cryo-ET (Atherton et al. 2022); if anything, our DCX-KO neurons show increased actin staining (Fig. S4C-D). In any case, our results on polyglutamylation do not exclude a link between DCX and actin. Indeed, given the complexity of MT-actin crosstalk (Dogterom and Koenderink 2019), it’s likely that the tubulin code impacts the actin cytoskeleton as well. We look forward to exploring these links in the future.

Sadly for us, our results argue against a role for DCX in the regulation of microtubule dynamics, which we and others had proposed based on DCX’s potent nucleation and anti-catastrophe activity in reconstitution assays (Moores et al. 2004; Bechstedt and Brouhard 2012). Rather, we only detected a minor misorientation of EB comets in our DCX-KO neurons (Fig. 3). The lack of change in EB comet behavior suggests that EB proteins outcompete DCX for their shared plus-end binding sites, e.g. via faster association kinetics (Reid et al. 2019), differences in binding site preferences (Ettinger et al. 2016), or exclusion of DCX from EB condensates (Song et al. 2023).

DCX also shares its binding site with its close relative, DCLK1. Both proteins regulate neuronal migration and morphology (Friocourt et al. 2007; Shin et al. 2013), and double-KO mice have severe defects in brain development (Deuel et al. 2006). Hence, DCLK1 might compensate for DCX-KO or interact differently with patient-mutant DCX. The kinase domain of DCLK1 is auto-regulatory (Agulto et al. 2021), effectively tuning the affinity of DCLK1 for microtubules, which may impact DCX’s affinity as well. Understanding the overlapping and independent roles of these two proteins is an important area of future research, including but not limited to their roles in the regulation of the tubulin code.

The PTMs of the tubulin code are frequently correlated; e.g., multiple PTMs are increased simultaneously by paclitaxel (Hammond et al. 2010), perhaps because stabilized microtubules have more time to accumulate PTMs. However, polyglutamylated microtubules specifically enhance detyrosination (Ebberink et al. 2022), indicating a more specific correlation, perhaps depending on the “lattice spacing” of the microtubule lattice (Yue et al. 2023). Our work demonstrates that one PTM can be significantly reduced (polyglutamylation) while others remain constant, indicating that PTMs can be individually regulated.

The identification of tubulin modifying enzymes has enabled significant progress in our understanding of the “tubulin code”, but we do not fully understand how these enzymes are regulated. Our work has identified DCX as a key regulator of polyglutamylation, and more generally implicates MAPs as potential regulators of tubulin PTMs. MAPs that regulate the tubulin code may share DCX’s sensitivity to structural features of the microtubule lattice. If the tubulin code has “writers” (like TTLLs) and “readers” (like spastin and tau), then DCX acts as a positive activator of a “writer”, somewhat like a good editor.

## Supporting information

Video S1_CTRL0 nuclear movements

Video S2_DCXKO1 nuclear movements

Video S3_CTRL0 SiRTubulin CellMaskActin

Video S4_CTRL0 SiRTubulin CellMaskActin

Video S5_CTRL0 SiRTubulin CellMaskActin

Video S6_DCXKO1 SiRTubulin CellMaskActin

Video S7_CTRL0 EB dynamics

Video S8_DCXKO1 EB dynamics

Video S9_CTRL0 Lysotracker

Video S10_DCXKO1 Lysotracker

Key resources table

## Data availability

Accession codes and unique identifiers for all biological reagents are provided in the Key Resource Table. All original data generated in this study is available upon request.

## Code availability

Custom code for nuclear and lysosome tracks analysis are available for download at https://github.com/hendricks-lab. Custom scripts for comet analysis and intensity profiles are available for download at http://sites.imagej.net/Hadim/.

## Acknowledgements

We would like to thank current members of the Brouhard, Hendricks and Bechstedt labs, for fruitful discussions during all phases of this project, and more particularly past members Hadrien Mary and Piper Stevens for their help in code writing. Many thanks to Geneviève Dorval and Thomas Durcan for thoughtful support with the use of iPSCs and CRISPR methods. We would also like to acknowledge the McGill University Advanced BioImaging Facility (ABIF), RRID: SCR_017697, the Flow cytometry core facility of the Life Science Complex, the Genome Québec sequencing platform, and their personal for providing services and helpful advice for data collection and analysis.

## Author contributions

Conceptualization and experimental design: GJB, MS, and AGH. Experimental procedures: MS and ALP, Data analysis: MS, ENNP and ALP, Manuscript writing: MS, GJB, and AGH. All authors participated in manuscript edition. Funding acquisition: GJB and AGH.

## Materials and Methods

### iPSCs origin, maintenance, and differentiation

Our use of stem cells was approved by McGill’s Institutional Review Board under permit number A12-M38-17A. hiPSCs initial stocks were obtained from the C-BIGR repository at the Montreal Neurological Institute. The AIW2-2 line comes from an unaffected male donor and was generated from PBMCs. The cells were grown on Matrigel coated surfaces in mTeSR plus media in feeder-free conditions, they were passaged using Gentle Cell dissociation reagent (GCDR) and supplemented with 10uM Y-27362 for 24h when required after thawing, gene-editing or low-density plating. hiPSCs were induced into Neuronal progenitor cells using Stemdiff Smadi induction media provided by Stemcell technologies. Briefly, the hiPSCs were seeded in a monolayer at high density and feeded with fresh induction media every other day for 6-12 days. Then the cells were dissociated using accutase and amplified on PO-laminin coated plates for an additional period of 10-15 days before cryopreservation or differentiation. Terminal differentiation into cortical neurons was performed using commercially available BrainPhys and supplemented as required by manufacturer’s guidelines. NPCs were seeded on PO-Laminin coated dishes or coverslips at variable densities depending on the experiment and were differentiated for 3 days. Cell fixation was performed following (Chazeau et al. 2016) methods: after 1,5min incubation in extraction buffer when specified (PIPES 80mM, EGTA 1mM, MgCl2 7mM, NaCl 150mM, D-Glucose 5mM, Triton X-100 0.3%, Glutaraldehyde 0.25%), PFA 4% warmed at 37°C was applied to the cells for 15min before rinsing and storage in PBS until further use.

### CRISPR gene-edition

AIW2-2 hiPSCs line was gene edited using CRISPR/Cas9 techniques. gRNAs for DCX Knock-In were designed using the CRISPRdirect online tool (https://crispr.dbcls.jp/). All the gRNA sequences used in this study are detailed in the key resource table. They were ordered as ssDNA oligos (IDT) and cloned into the pX458 plasmid generously gifted by Feng Zhang (#48138), using the BBsI restriction site. The newly generated plasmids were transformed into Stbl3 strains and amplified according to nucleobond Maxiprep kit supplier’s recommendations. The actual gene edition was performed using electroporation, as follows: the hiPSCs were detached from the substrate using accutase, counted and about 5millions cells were resuspended in 100ul of cold Internal Neuronal Buffer (KCl 135mM, CaCl2 200µM, MgCl2 2mM, HEPES 10mM, EGTA 5mM). The cell suspension was added to 5ug of gRNA/Cas9 plasmid with 30ug of the corresponding single stranded template before application to a 100ul BTX cuvette. The BTX Gemini X2 twinwave electroporator was used to deliver a single 10ms pulse at 60V. Warmed up mTeSR was then added on top of the electroporated cells before transfer of the cell suspension to a new matrigel coated dish with rock inhibitor supplemented mTeSR. Three days post-electroporation, the cells were again dissociated from the substrate using accutase and were passed through a 70um strainer before FACs-sorting for GFP positive cells. The cell sorting was performed in the Flow Cytometry Core Facility for flow cytometry and single cell analysis of the Life Science Complex and supported by funding from the Canadian Foundation for Innovation. The positive cells were seeded in individual wells of 96 well plates and left to grow for about 2 weeks. For proper recovery and growth of the cells, rock inhibitor or CloneR were kept in the mTeSR media for 24 to 72h post-sorting. After sufficient growth of individual hiPSCs clones in the 96 well plates, the clones were collected and split in two for amplification and screening for indels in extracted genomic DNA. For screening, each clone’s genomic DNA was extracted in Quick extract buffer, amplified by PCR (see key resource table for primer sequences) and depending on the profile of the PCR product and response to T7 assay, the clones were subjected to Sanger sequencing performed at Genome Québec. The gene edited iPSCs were tested for presence of pluripotency markers (OCT4, SSEA3), amplified, and cryopreserved in FBS-20%DMSO until further use for differentiation.

### Neuronal cell preparation

Following fixation, the coverslips were quenched in NaBH4 at 1mg/ml for 10min, then they were subjected to permeabilization in PBS-Triton 0.5% for 10min. Unspecific bindings were blocked by bathing the coverslips in PBS-Triton0.1%-BSA3% for 30 min before incubation with the primary antibodies (see key resource datasheet) overnight at 4°C. Incubation with the secondary antibodies were done at room temperature for 1h, before a DAPI stain and mounting of the coverslips with Fluorsave.

Transient expression of EB3-mCherry, DCX-GFP or TTLL11-mRuby in neurons were performed using electroporation. Briefly, the NPCs were dissociated and about 0.5-1 million cells were mixed with 5ug of plasmid in 200ul of INB. The electroporation was performed with a 1ms pulse at 350V in 620 BTX cuvettes. The cells were then seeded on glass bottom dishes or coverslips and imaged in BrainPhys or fixed as previously described. When dyes were used to look at specific structures in live neurons, they were all diluted in warmed up BrainPhys and incubated for specified timing with the cells before removal and imaging: SiR tubulin 100nM 30min, CellMask Green Actin 0.5X 30min, Lysotracker 50nM 10min, Syto Nucleus Deep red 250nM 30min. For sorting of TTLL11 electroporated neurons, electroporation was performed similarly as described above, with 10 million cells and 12 µg of plasmid. After electroporation, the cells were seeded on regular plastic dishes and cultured in terminal differentiation media (BrainPhys) for 2 days. On the second day, the cells were collected using accutase, sorted at the FACs sorting facility based on their level of mRuby fluorescence, and subsequently plated again at approximately 15000 cells/cm^2^ in BrainPhys media for one additional day. The cells were then fixed and stained for polyE and Tubb3 as described above.

### TTLL11 Plasmid generation

The human TTLL11 primary transcript (NCBI: NM_001139442) in frame with restriction sites at the N and C-terminal ends of the sequence, was synthesized by IDT after manual codon optimization (see key resource table for sequence). The gene was digested by HindIII and AgeI enzymes alongside a pCMV-mRuby plasmid generously gifted by Pr. Arnold Hayer. The digested TTLL11 gene was then ligated into the linearized mRuby plasmid. The validity of this construct was assessed by sequencing and expression in model cells. Moreover, polyglutamylation levels increased significantly upon expression, confirming the polyglutamylase activity of the expressed TTLL11.

### Cell line culture, transfection

Human osteosarcoma cells (U2OS) were used as a secondary non-neuronal model system. The cells were grown at 37°C and 5% CO2 in high glucose DMEM supplemented with 10% serum and 1% antibiotic. Once seeded at 40000 cells/cm^2^ on coverslips, the cells were transfected with an equivalent amount of DNA: either 1.5µg of TTLL11-mRuby, 1.5µg of DCX-GFP, or 1.5µg total of both plasmids, using the calcium phosphate method [Kingston 2003]. The cells were left to express 24h before extraction (PIPES 80mM, EGTA 1mM, MgCl2 7mM, NaCl 150mM, D-Glucose 5mM, Triton X-100 0.3%, Glutaraldehyde 0.25%) for 1min30s and fixation for 10min using cold Methanol, or lysis (2X SDS loading buffer: 125mM Tris-HCl pH 6.8, 20% glycerol, 0.04% Bromophenol blue, 4% SDS, 100mM DTT). The coverslips were then stained for polyE as described above.

### Western blots

The cells were lysed by incubation in cold RIPA or 2X SDS loading buffer (125mM Tris-HCl pH 6.8, 20% glycerol, 0.04% Bromophenol blue, 4% SDS, 100mM DTT) and were collected by scraping the plates. Lysates in RIPA buffer were supplemented with complete-T for long term storage. Before loading onto a 12% acrylamide gel (precasted, Genscript), the lysates in RIPA were mixed with 2X SDS loading buffer and all samples were sonicated, spun down and boiled for 10min at 95°C. Electrophoresis was done in MOPS buffer at 160 constant voltage, then proteins were transferred to nitrocellulose membranes and blocked in 5% Milk. Primary antibodies were diluted according to specific dilutions (see key resource datasheet) and incubated with the membrane at 4°C overnight. The membranes were rinsed with TBST and incubated for 2h with anti-HRP secondary antibodies. The proteins were revealed using ECL pico, and 2s-30min membrane exposure to Diamed films. Since DCX, native tubulin and modified tubulin have similar molecular weight, all films are from different loadings-gels of the same samples, at the exception of the separation of α and β-tubulin where SDS-PAGE gels were prepared and used as described in [Banerjee 2010].

### Imaging and analysis

Image collection for this manuscript was performed in the McGill University Advanced BioImaging Facility (ABIF), RRID:SCR_017697. All images were analyzed primarily in Fiji and processed using Microsoft office excel, and python. The Zeiss LSM710 inverted confocal and X-Cite 120 LED with Zen 2.0 software were used to screen and acquire images of fixed samples. The objectives 10X EC plan neofluar NA 0.3, 20X plan apochromat NA 0.8 and 63X plan apochromat NA 1.4 oil were used. Intensity profiles in neuronal fixed samples were obtained from thick ROI lines drawn on neurites from the tip to the cell body, or by creation of a dilated mask of the DAPI signal for cell bodies. The intensity per pixel was interpolated from pixels along the width of the line, the generated profiles were then further compiled with other images using excel and python. For intensity quantification in entire field of views, all images were acquired in the same laser power and gain conditions, the Tubb3 or tyrosinated (YL1/2) microtubule signal was used as a mask for cells and intensity with the other channels were measured inside the mask and normalized to the mask intensity. The number of branches per cell were manually counted and their length measured with fiji line tools. For DCX and TTLL11 expression in U2OS cells, ROIs were drawn around the expressing cells and signal intensity for DCX-GFP, TTLL11-mRuby and polyE (Alexa-647) were measured and normalized to the cell area.

The Quorum WaveFX-X1 spinning disk confocal system on a Leica DMI6000B inverted microscope equipped with a Photometrics prime BSI Cmos camera was used for short term live-cell imaging. We used the 63X plan apochromat NA 1.4 oil objective and the Metamorph software to record EB3mCherry and lysotracker dynamics. All timelapse movies recorded were 2min total, and live cells were never imaged for more than 1h. For EB3, images were taken with 2s interval, in neurons with low-medium expression levels. Kymographs were done from wide multipoint ROI lines drawn on processes from the tip to the cell body. Then straight ROI lines were drawn on each comet and a homemade python script extracted the comet direction, lifetime, length and growth rate. For each cell, the growth cone and process areas were measured by thresholding a maximum projection of the timelapse. All further analyses were made in python and used a mean value per cell.

For lysotracker data acquisition, the timelapses were 500ms interval. Straight sections of the processes excluding cell bodies were selected using the polygon ROI tool and cargo trajectories were analysed inside, using the Trackmate plugin (Ershov et al. 2022; Tinevez et al. 2017). Trackmate was set up to use the LoG detector to identify 1µm diameter spots, with subpixel localization. Then the simple LAP tracker was used to define and filter trajectories that were exported as xml files. To analyze the TrackMate data, we had previously written custom MATLAB codes (Prowse et al. 2023), available on Github: https://github.com/hendricks-lab. For each trajectory, we converted coordinate x and y position values to a single position vector with respect to the location of the soma. This position vector had a positive value when the tracked cargo moved anterogradely, and a negative value when moving retrogradely, as shown in Figure 4. Using the position vector, we reduced the noise using a Savitzky-Golay smoothing filter of power 2 and span of 9 and segmented the trajectory into runs at each inflection point. These runs were used in the run length and velocity analysis. Between each inflection point, we determined the absolute value of the cargo’s displacement. If the displacement within the run was larger than 2 μm, the run was considered processive, while if it was between 0.03 μm and 2 μm, the run was categorized as diffusive, and finally if the displacement was 0.03 μm or smaller, the run was considered stationary. The threshold for a run to be considered processive was determined by plotting the α value of processive runs compared to that of diffusive runs, as well as the directional bias of diffusive and processive runs. We determined that 2 μm was optimal because the processive runs had an α value near 2, while the diffusive runs had an α near 1, as expected. We also observed that there was no directional bias for diffusive runs at the set minimum run length threshold, indicating the runs are purely diffusive. We then determined the run length and velocity per trajectory by taking the mean of the processive runs or processive runs over time, respectively, for each trajectory. We calculated the α by calculating the slope of the mean squared displacement plot for each trajectory and subsequently taking the mean for each cell. We calculated mean squared displacement using a time average for time intervals from the exposure time of 0.5 s up to 20 s. We used (Eq. 1) to calculate the mean squared displacement (MSD), where t is the frame, τ is the time interval, and x is the position vector value at the specified time.

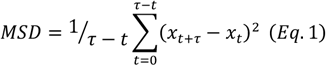

The first 5 s of log-log plot of the mean squared displacement is used to calculate the α value for each trajectory, to ensure only the linear portion is fitted. We calculated the radius of gyration (Rg) using (Eq. 2). Here, r represents the sum of the squared x and y components, and N represents the number of position values in the entire trajectory.

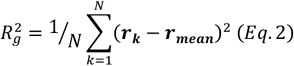

To determine the directional bias, we determined the fraction of time the processive runs spent moving anterogradely compared to retrogradely, using the identified processive runs from the previously described function analyzing run length and velocity. To calculate the fraction of processive, stationary, and diffusive runs, we calculated the fraction of time for each run within each trajectory spent in each of these states, as defined by the previously described thresholds.

The perkin-elmer high content screening microscopes operaphenix in confocal mode with a 20X NA 1.0 water objective and the Harmony 9.0 software were used for long time recording of neuronal movements and branch dynamics. Images were taken at 5-10min interval for 8 to 15h. For each imaging session, multiple FOV from the same well were taken and analysed individually. Nuclei tracking was done using trackmate within an automated python-based pipeline. The LoG detector and LAP simple tracker were used with subpixel detection of 10µm spots, and 40µm maximum linking distance. Trajectories and spot data were exported as xml files, visually checked, and compiled within python. Only the trajectories that lasted more than 30min in total were included in the analysis.

### Statistical analysis

All statistical tests used are indicated in the figure legends: Kruskal-Wallis non parametric H test, Mann-Whitney non parametric U test, or Student’s t test. Pvalues were considered nonsignificant when above 0.05.

For the lysotracker dataset: For each per trajectory analysis, we took 1000 bootstrapped means with replacement and averaged them the same number of times as the minimum number of trajectories from either dataset. We then compared these means by subtracting the DCX-KO data from the control data and comparing the fraction of the difference distribution overlapping with 0. If this fraction was less than 0.05%, then we considered it a statistically significant difference, with a p value equal to the fraction that overlaps with the 0 line. Each value calculated per cell was analyzed using Student’s two-tailed t test.

**Figure S1:**
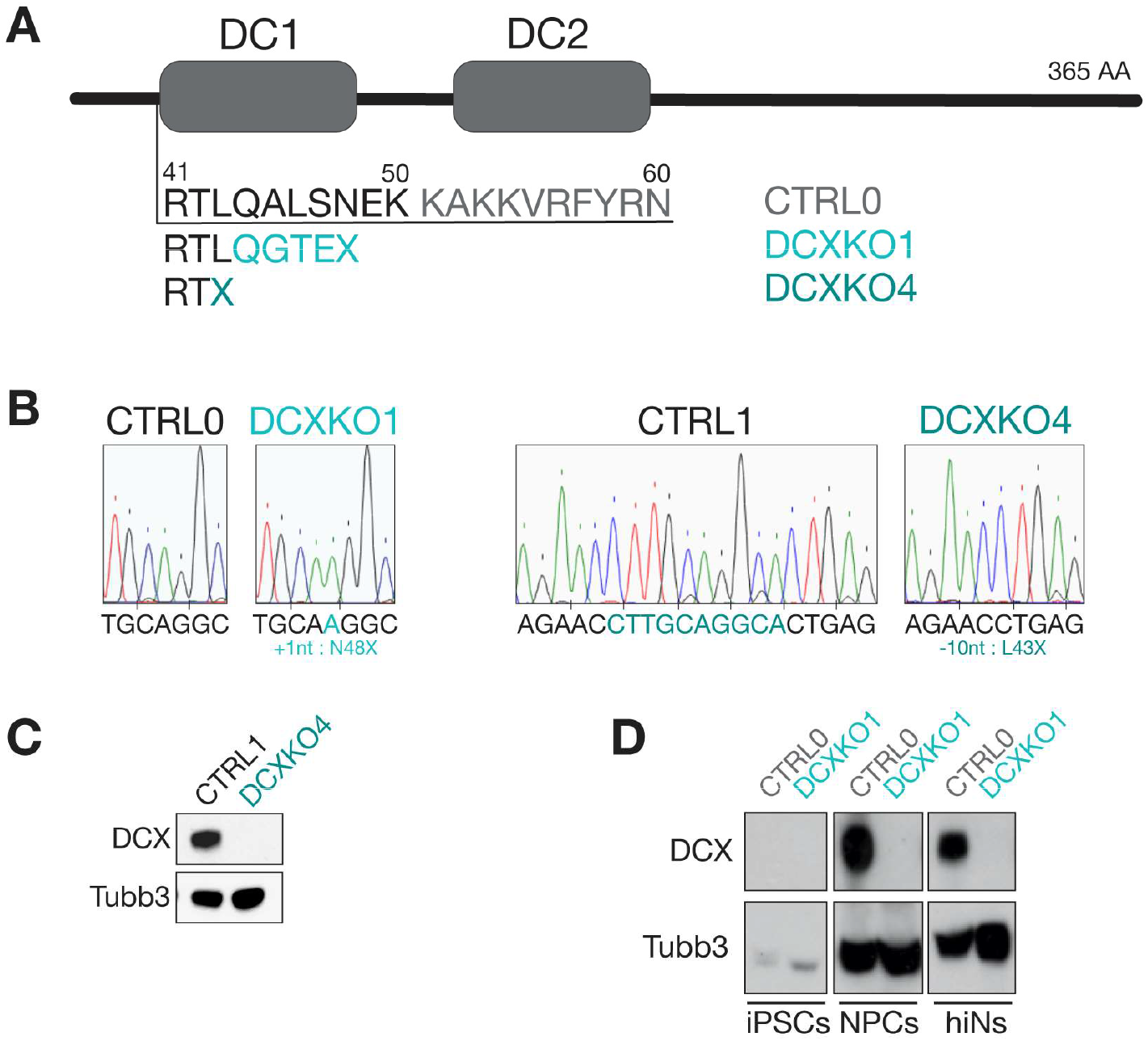
Engineering and sequences DCXKO1 and DCXKO4. **(A)** DCX contains two microtubule binding domains, DC1 and DC2. DCX-KO cells were obtained by indels introduced in *Dcx*’s gene sequence before DC1. CTRL0/CTRL1, DCXKO1 and DCXKO4 amino acid sequences are shown from residue 41 to 60. The asterisk shows the indel location and the resulting amino acid sequence. **(B)** Chromatograms showing nucleotide sequences for CTRL0, DCXKO1, CTRL1 and DCXKO4, with indels highlighted below. **(C)** Immunoblot of CTRL1 and DCXKO4 neuron lysates at DIV3, with antibodies against Tubulin-β3 (Tubb3) and DCX. **(D)** Immunoblots of CTRL0 and DCXKO1 lysates at the iPSCs stage, the NPCs stage and the neuronal stage at DIV3, with antibodies against Tubb3 and DCX. The same samples were loaded on different gels for both antibodies.

**Figure S2:**
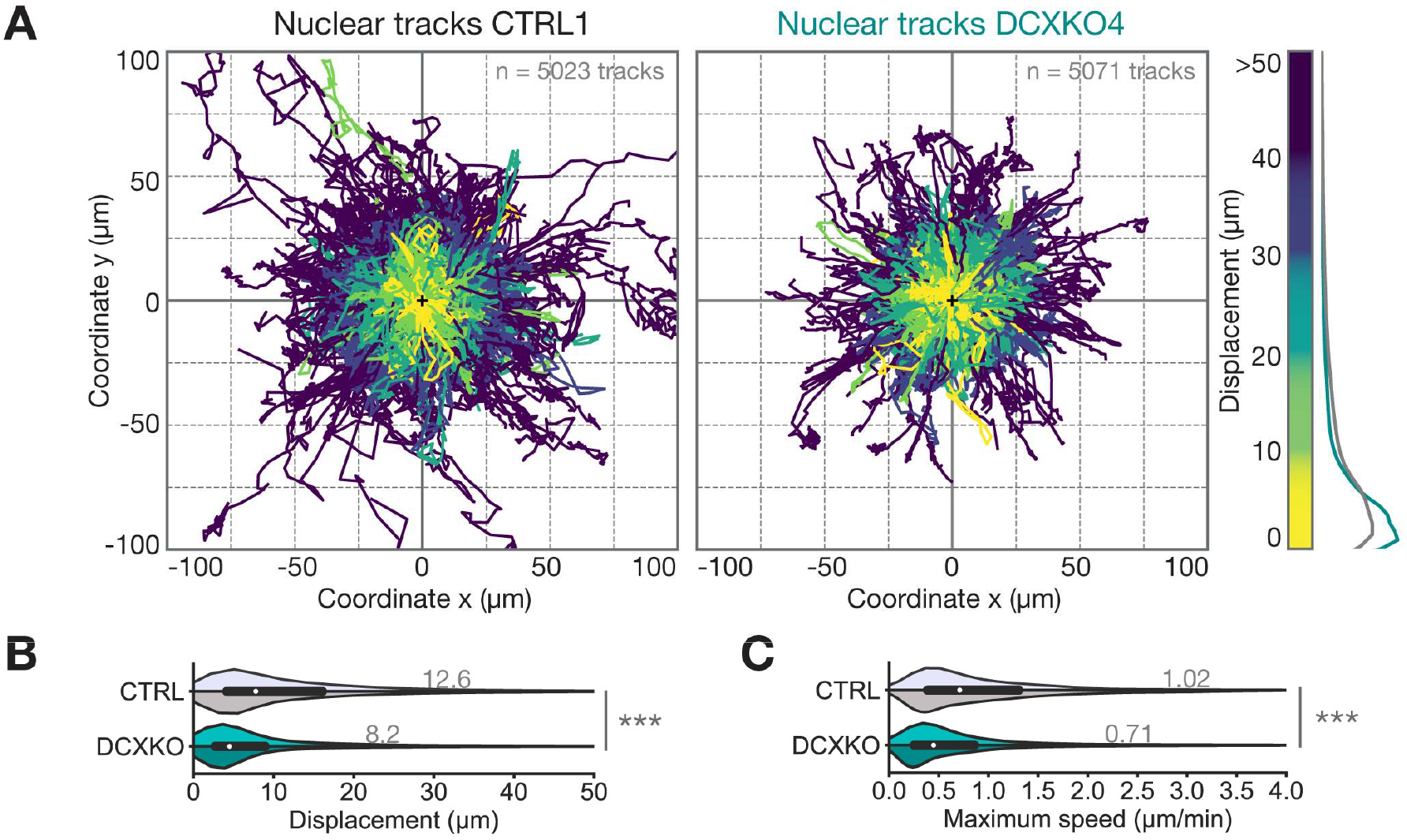
DCXKO1 and DCXKO4 neurons experience reduced migration speed and efficiency. **(A)** Position plots showing a sample of trajectories extracted from CTRL1 (left) and DCXKO4 (right) nuclei tracking. All nuclei are considered to be at the same coordinates (0,0) at their first timepoint. Tracks are color-coded according to the displacement of the nucleus. Distribution of displacement for both genotypes are shown on the right end side of the panel. *n*=5023 and 5071 nuclei respectively for CTRL1 and DCXKO4. **(B)** Distribution of displacement achieved by CTRL0 (light grey), CTRL1 (dark grey), DCXKO1 (cyan) and DCXKO4 (dark cyan) nuclei during the imaging period. **(C)** Distribution of maximum speed achieved by CTRL0 (light grey), CTRL1 (dark grey), DCXKO1 (cyan) and DCXKO4 (dark cyan) nuclei during the imaging period. For B-C, *n*=21826 tracks for CTRL0, 24801 tracks for CTRL1, 6461 tracks for DCXKO1 and 14837 tracks for DCXKO4. Each line was used in 3 paired experiments, tracks with less than 30min of imaging time were excluded from the dataset. These two plots were cut-off to better show the main population, but the maximum values for displacement and max speed are 287.07 µm, and 13.22 µm/min respectively. Statistical analysis was performed using the Kruskal-Wallis non parametric H test: *** *p*<0.001.

**Figure S3:**
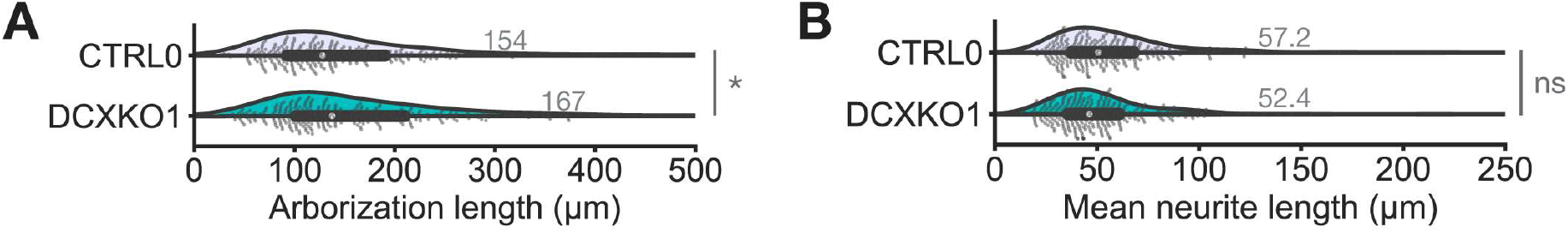
DCXKO1 and DCXKO4 have similar mean neurite length at DIV3. **(A)** Distribution of arborization length for fixed CTRL0 and DCXKO1 neurons used in the neurite counts. **(B)** Distribution of mean neurite length for fixed CTRL0 and DCXKO1 neurons used in the neurite counts. For A-B *n*=323 control and 351 DCXKO1 neurons from 3 experiments. These two plots were cut-off to better show the main population, but the maximum values for arborization length and mean neurite length are 912.27 µm and 342.78 µm respectively. Statistical analysis was performed using a Mann-Whitney U test: ns nonsignificant, * *p*<0.05.

**Figure S4:**
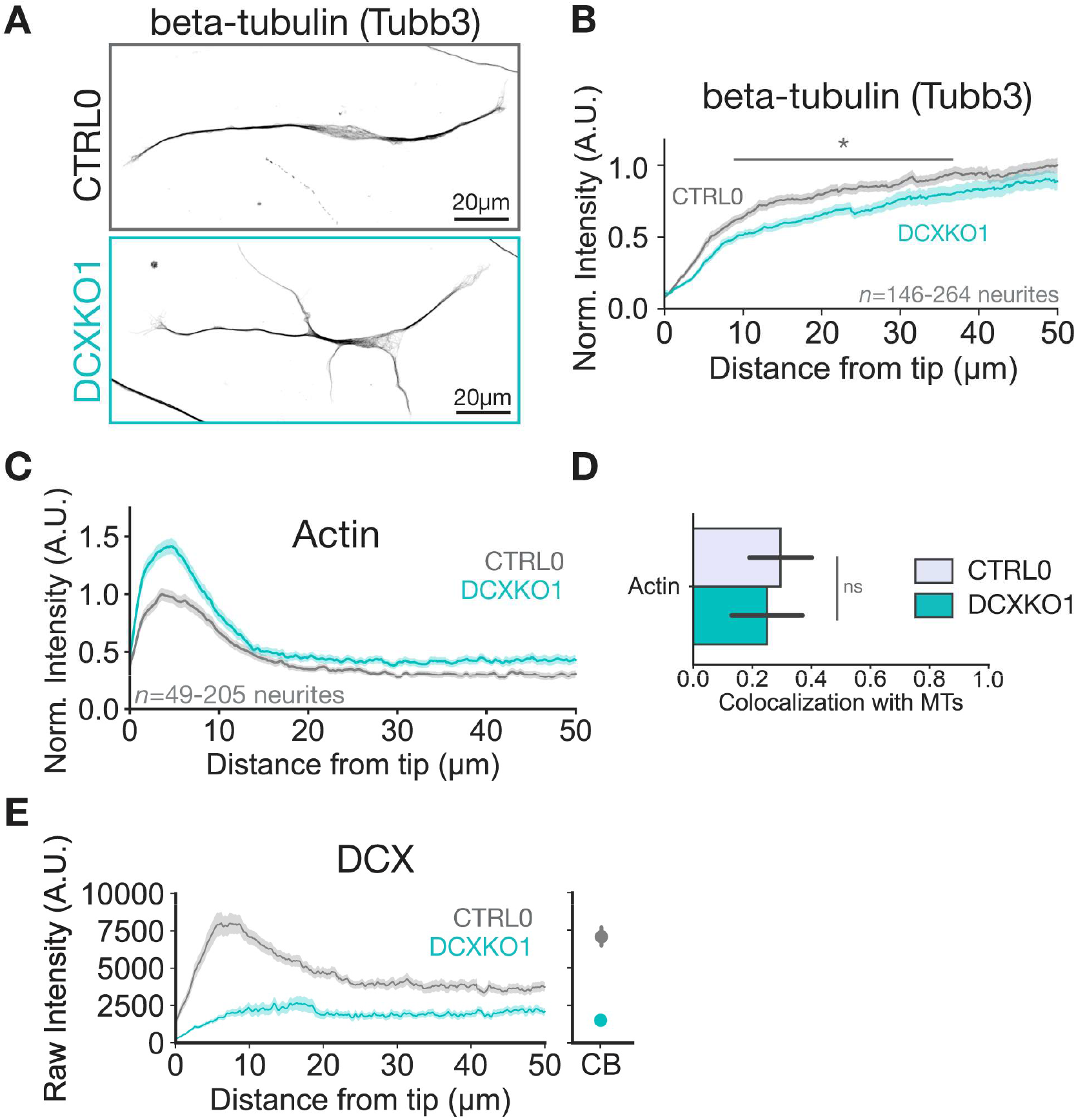
Modifications to tubulin and actin network in DCXKO neurons. **(A)** Representative immunostaining of CTRL0 and DCXKO1 neurons at DIV3 with an antibody against β3-Tubulin (Tubb3). **(B)** Mean β-Tubulin intensity profile +/- SEM. found in the distal tip of neurons for CTRL0 and DCXKO1. Images included in the analysis are from *n*=146 CTRL0 neurites and *n*=264 DCXKO1 neurites from 4 experiments. Statistical analysis was performed using the Mann-Whitney non parametric U test: ns nonsignificant. **(C)** Mean Actin intensity profiles +/- SEM. found in the distal tip of neurons for CTRL0 and DCXKO1. Images included in the analysis are from *n*=205 CTRL0 and *n*=145 DCXKO1 neurites from 3 independent experiments. **(D)** Percentage colocalization of Phalloidin with Tubb3 in the GC of neurons. *n*=65 CTRL0 GC and *n*=159 GC neurites from 2 experiments. Statistical analysis was performed using the Mann-Whitney non parametric U test: ns nonsignificant. **(E)** Raw DCX intensity profiles +/- SEM. found in the distal tip and cell body (CB) of neurons for CTRL0 and DCXKO1. Images included in the analysis are from *n*=127 CTRL0 and *n*=49 DCXKO1 neurites and n=306 CTRL0 and 365 DCXKO1 cell bodies from 2 independent experiments.

**Figure S5:**
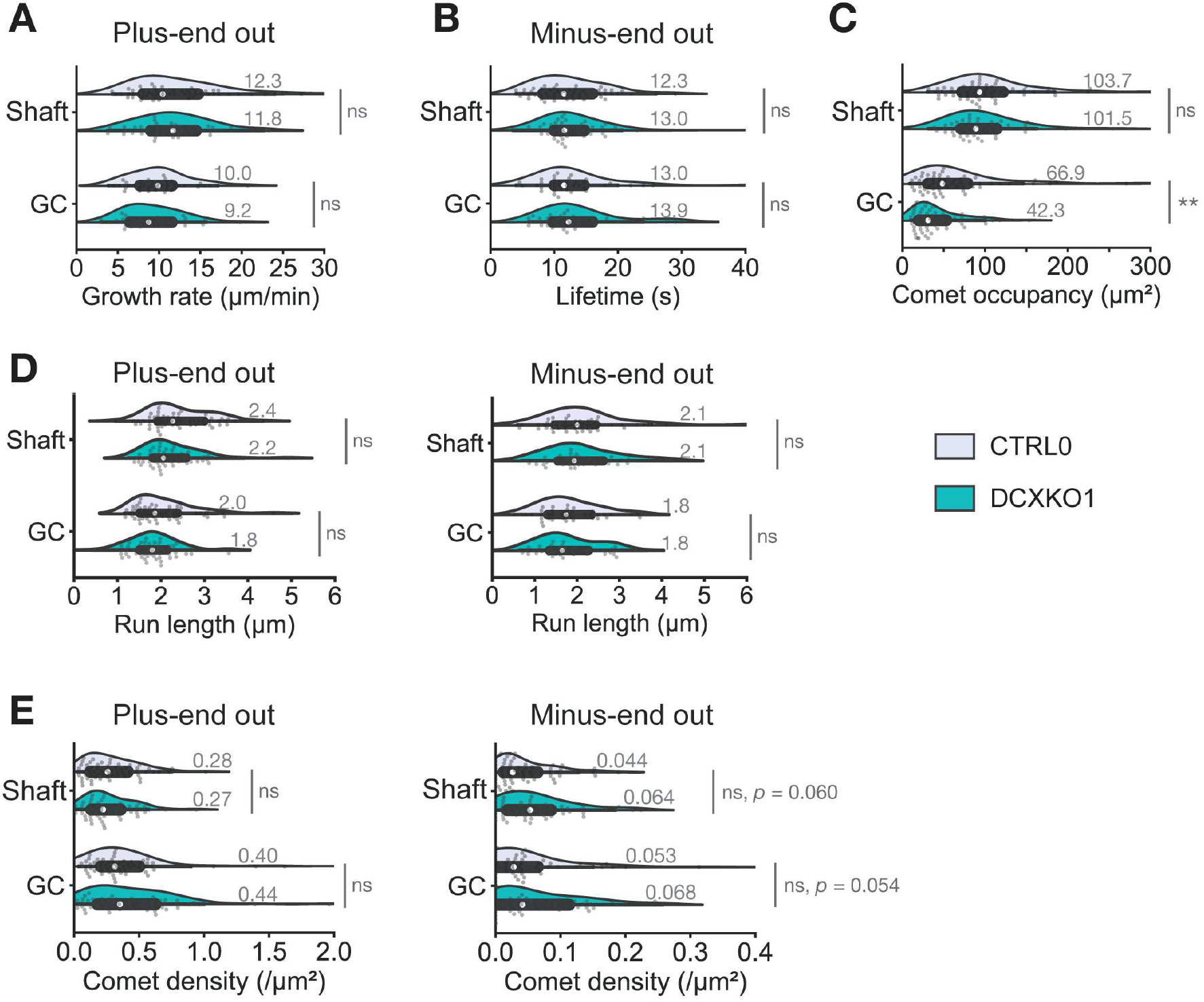
Microtubule dynamics are not different in DCXKO1 neurons. **(A)** Violin plot showing the distribution of growth rates for minus ends out comets in shaft or GC of CTRL0 and DCXKO1 neurons. **(B)** Violin plot showing the distribution of lifetimes for minus ends out comets in shaft or GC of CTRL0 and DCXKO1 neurons. **(C)** Violin plot showing the comet occupancy or area covered by EB3 comets during the imaging period. **(D)** Violin plot showing the distribution of run length for plus-end out and minus-end out comets in shaft or GC of CTRL0 and DCXKO1 neurons. **(E)** Violin plot showing the distribution of comet density for plus-end out and minus-end out comets in shaft or GC of CTRL0 and DCXKO1 neurons. Some plots were cut-off to better show the main population, but the maximum values for comet density and comet occupancy are 2.14 µm^-1^ and 438.10 µm^2^ respectively. For A-E, *n*=63 control and *n*=63 DCXKO cells from 4 experiments. Statistical analysis was performed using the Mann-Whitney non parametric U test: ns nonsignificant, ** *p*<0.01.

**Figure S6:**
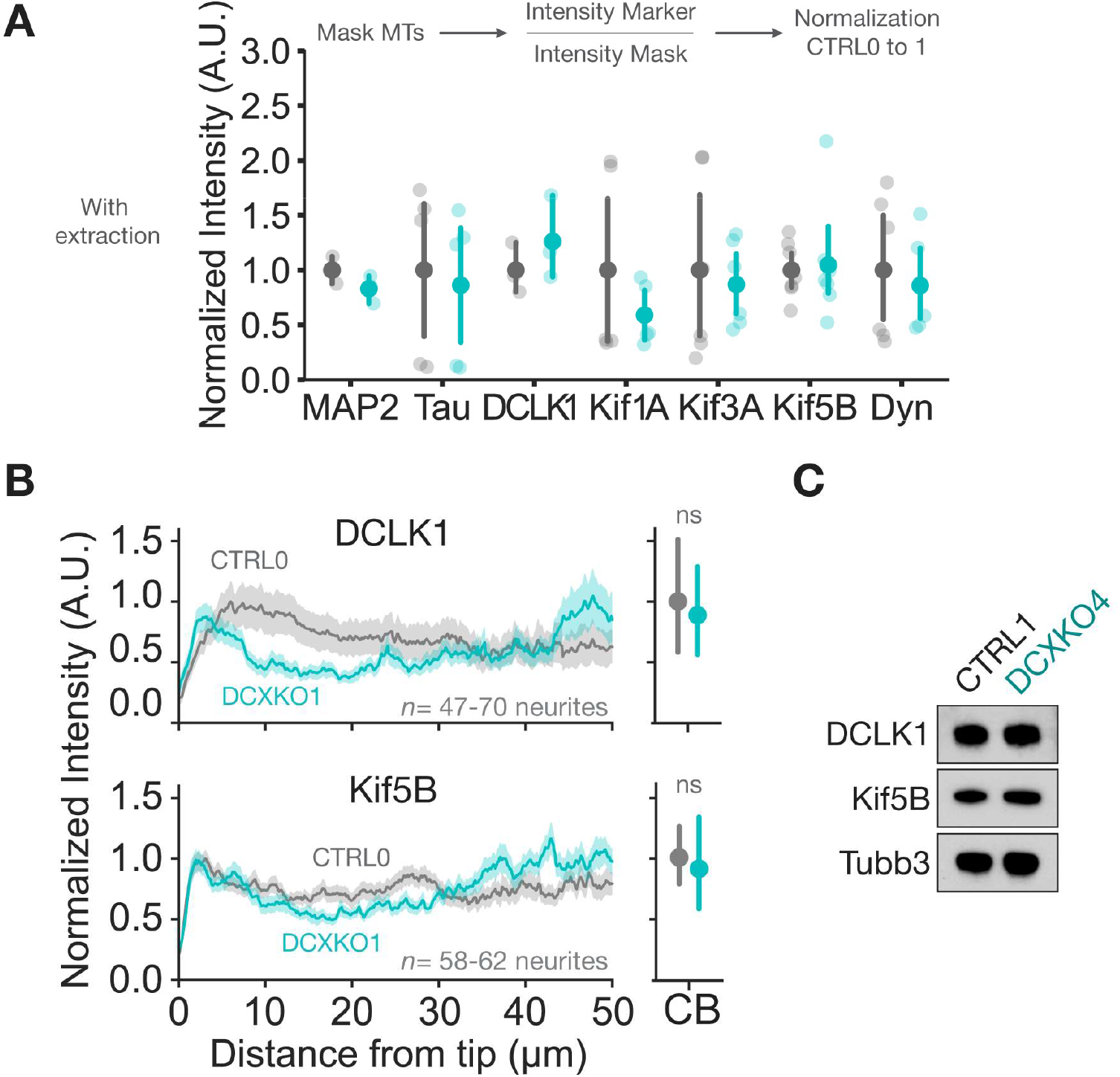
DCLK1 and Kif5B levels in DCX-KO neurons. **(A)** Compared mean intensity for MAP2, Tau, DCLK1, Kif1A, Kif3A, Kif5B, and Dynein in CTRL0 or DCXKO1 DIV3 extracted and immuno-stained neurons. The analysis pipeline is shown above the plot: a mask of the microtubules (MTs) was made, and the intensity of the marker was normalized to the intensity of the mask and to the control condition. Each transparent point represents a single image, while the opaque points show mean values, error bars represent the confidence intervals. Analyses are from *n*=3-8 CTRL0 or DCXKO1 images from 2-4 experiments. Statistical analysis was performed using the Mann-Whitney non parametric U test: ns nonsignificant. **(B)** Mean intensity profiles +/- SEM. of DCLK1 and Kif5B neurites distal ends and cell bodies (CB) +/- confidence interval for CTRL0 and DCXKO1 neurons. Images included in the analysis are from *n*=47-58 CTRL0 and *n*=70-62 DCXKO1 neurites for DCLK1 and Kif5B respectively from 4 experiments. Statistical analysis was performed using the Mann-Whitney non parametric U test: ns nonsignificant. **(C)** Immunoblots of CTRL1 and DCXKO4 neuron lysates at DIV3, with antibodies against DCLK1, Kif5B and Tubb3. The same samples were loaded on different gels for each antibody.

**Figure S7:**
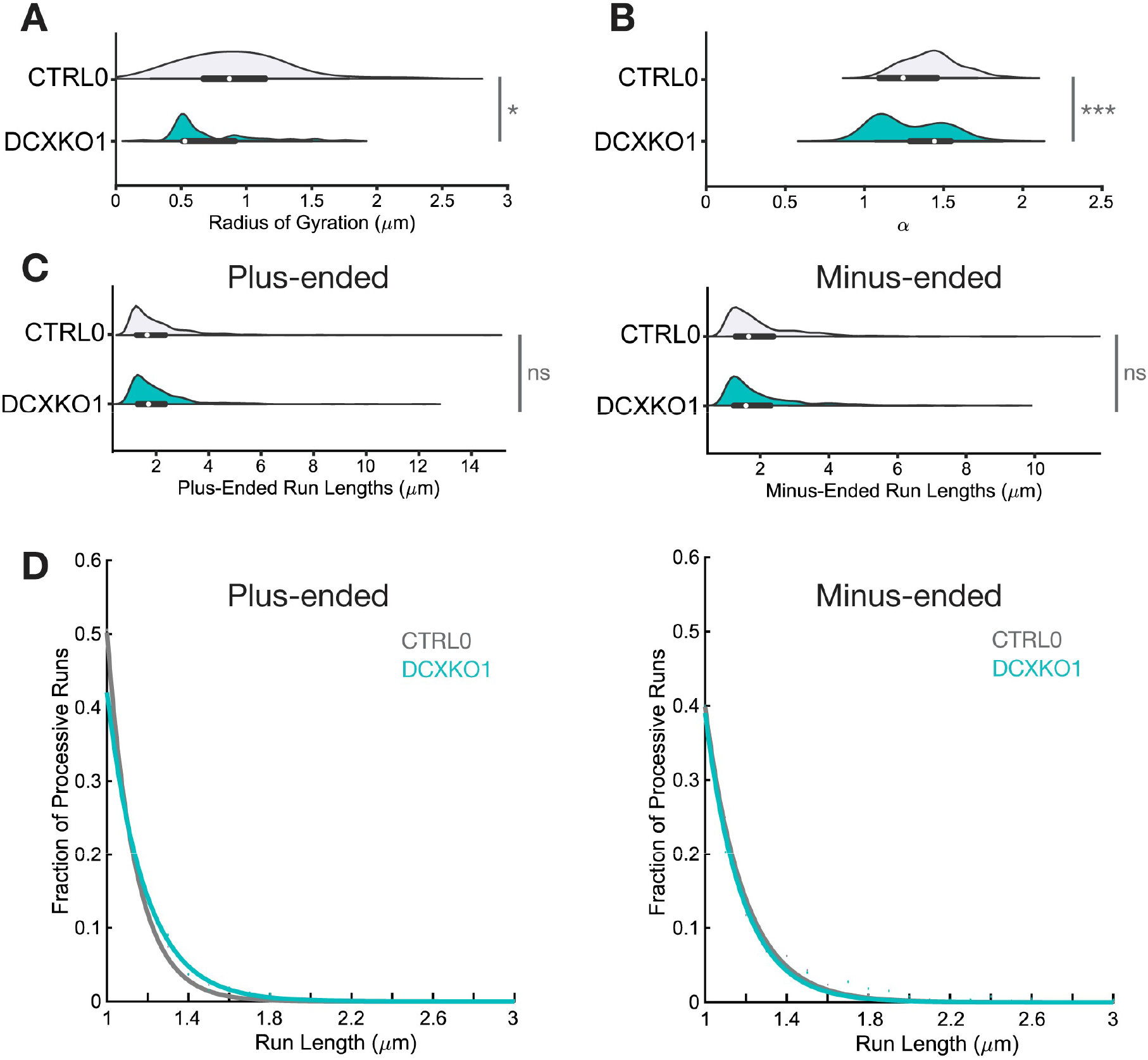
Alpha values and Rg for Lysosomes in DCXKO1. **(A)** Violin plot showing the radius of gyration of lysosomes in CTRL0 and DCXKO1 neurons. **(B)** Violin plot showing the alpha value of lysosomes in CTRL0 and DCXKO1 neurons. **(C)** Violin plot showing the distribution of plus-ended and minus-ended run length of lysosomes in CTRL0 and DCXKO1 neurons. **(D)** Fraction of plus-ended and minus-ended processive runs of lysosomes in CTRL0 and DCXKO1 neurons, depending on their run lengths. All trajectories (6082 for control and 3887 for DCXKO1) were obtained from *n*=41 control and 30 DCXKO1 neurons from 3 independent experiments. Statistical analysis was performed using the Student’s t-test: ns nonsignificant, * *p*<0.05, *** *p*<0.001.

**Figure S8:**
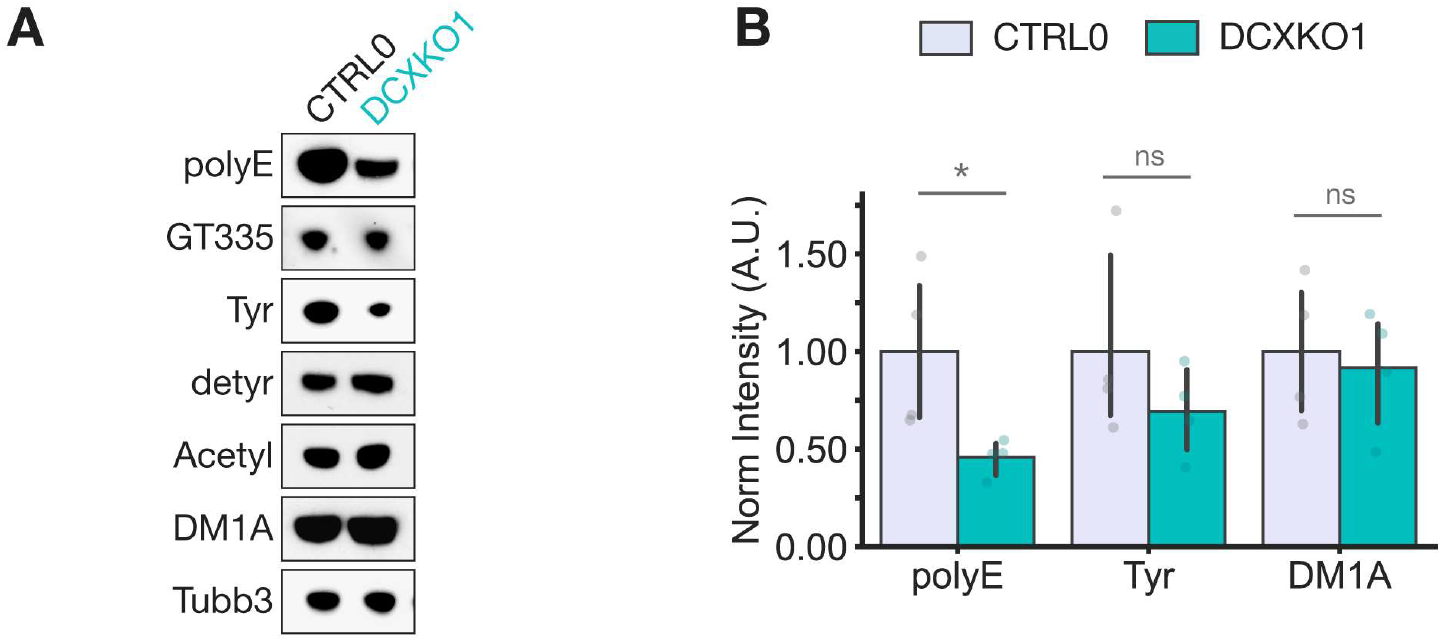
PTMs levels in DCXKO neurons. **(A)** Immunoblots showing the relative post-translational modifications levels in CTRL0 and DCXKO1 DIV3 neurons. The same samples were loaded on different gels for each antibody. **(B)** Immunoblot quantification showing polyglutamylation (polyE), tyrosination (Tyr) and α-tubulin (DM1A) levels normalized to tubulin-β3 (Tubb3) in CTRL0 and DCXKO1. *n*=4 CTRL0 and 4 DCXKO1 independent lysates. Statistical analysis was performed using the Student’s t-test: ns nonsignificant, * *p*<0.05.

**Figure S9:**
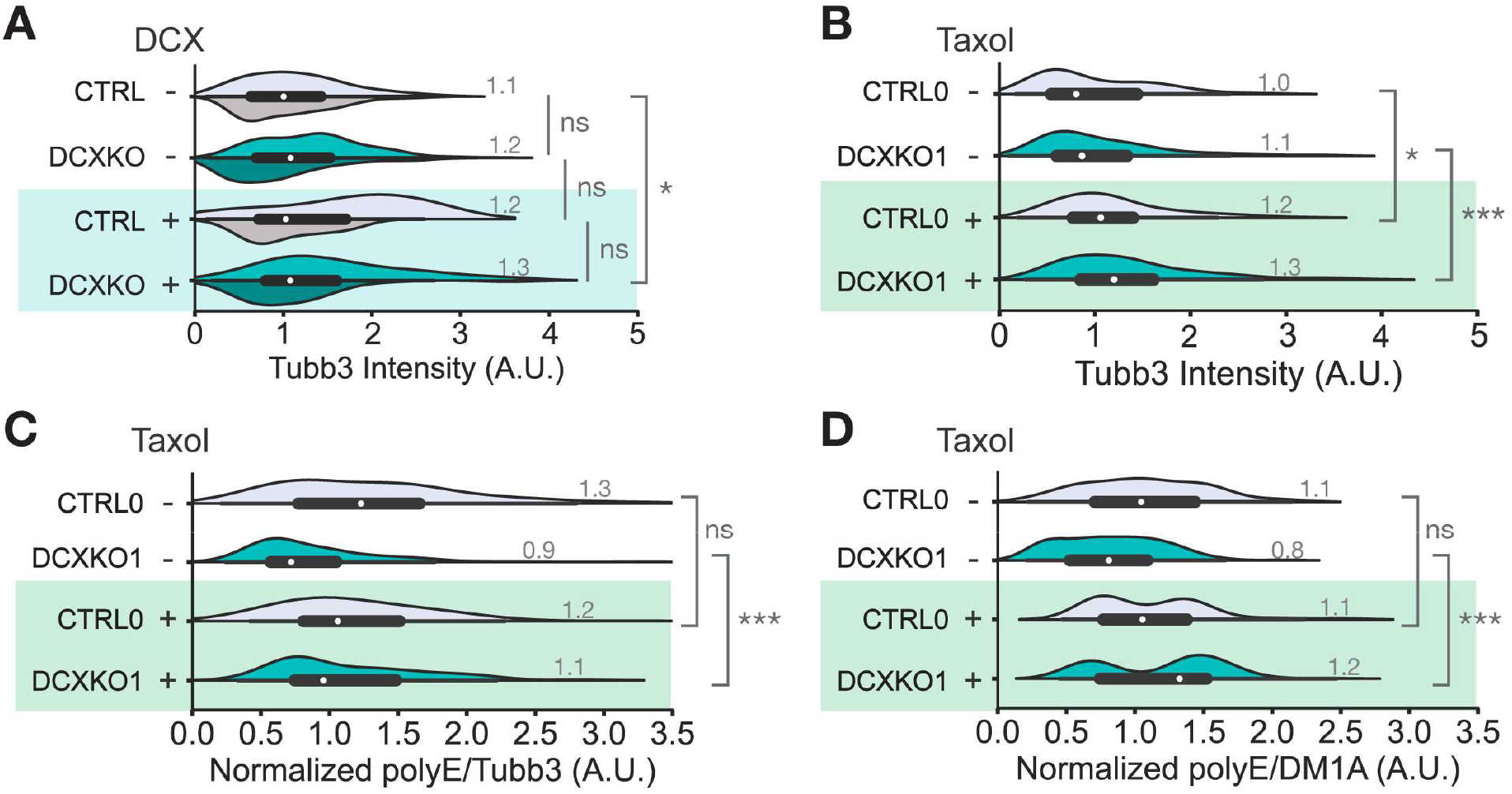
DCX reduces neurite numbers by increasing tubulin polyglutamylation levels. **(A)** Violin plots showing the Tubb3 intensity in CTRL and DCXKO DIV3 neurons with or without DCX-GFP electroporation. Images included in the analysis are from *n*=97 CTRL0 (light grey), 178 CTRL1 (dark grey), 225 DCXKO1 (cyan), 171 DCXKO4 (dark cyan), 11 CTRL0+DCX-GFP, 34 CTRL1+DCX-GFP, 38 DCXKO1+DCX-GFP, and 45 DCXKO4+DCX-GFP neurons. Each line was used in 3 paired experiments. Statistical analysis was performed using the Mann-Whitney non parametric U test: ns nonsignificant, * *p*<0.05. **(B)** Violin plots showing the Tubb3 intensity in CTRL0 and DCXKO1 DIV3 neurons with or without 100nM Taxol for 48h. **(C)** Violin plots showing the polyglutamylation (polyE) intensity in CTRL0 and DCXKO1 DIV3 neurons with or without 48h 100nM Taxol treatment (Tx), normalized to Tubb3 intensity. This plots was cut-off to better show the main population, but the maximum value for polyE/Tubb3 intensity is 7.12 (A.U.). **(D)** Violin plots showing the polyglutamylation (polyE) intensity in CTRL0 and DCXKO1 DIV3 neurons with or without Taxol treatment (Tx), normalized to DM1A intensity. For B-D, images included in the analysis are from *n*=84 CTRL0, 172 DCXKO1, 120 CTRL0+taxol, and 160 DCXKO1+taxol neurons from 2 experiments. Statistical analysis was performed using the Mann-Whitney non parametric U test: ns nonsignificant, * *p*<0.05, *** *p*<0.001. To note: as taxol treatment selectively enhances Tubb3 intensity, the effect of the treatment on polyE/Tubb3 seem lessen. However, taxol rescues polyglutamylation levels in DCXKO1 independently of the normalization method used.

**Figure S10:**
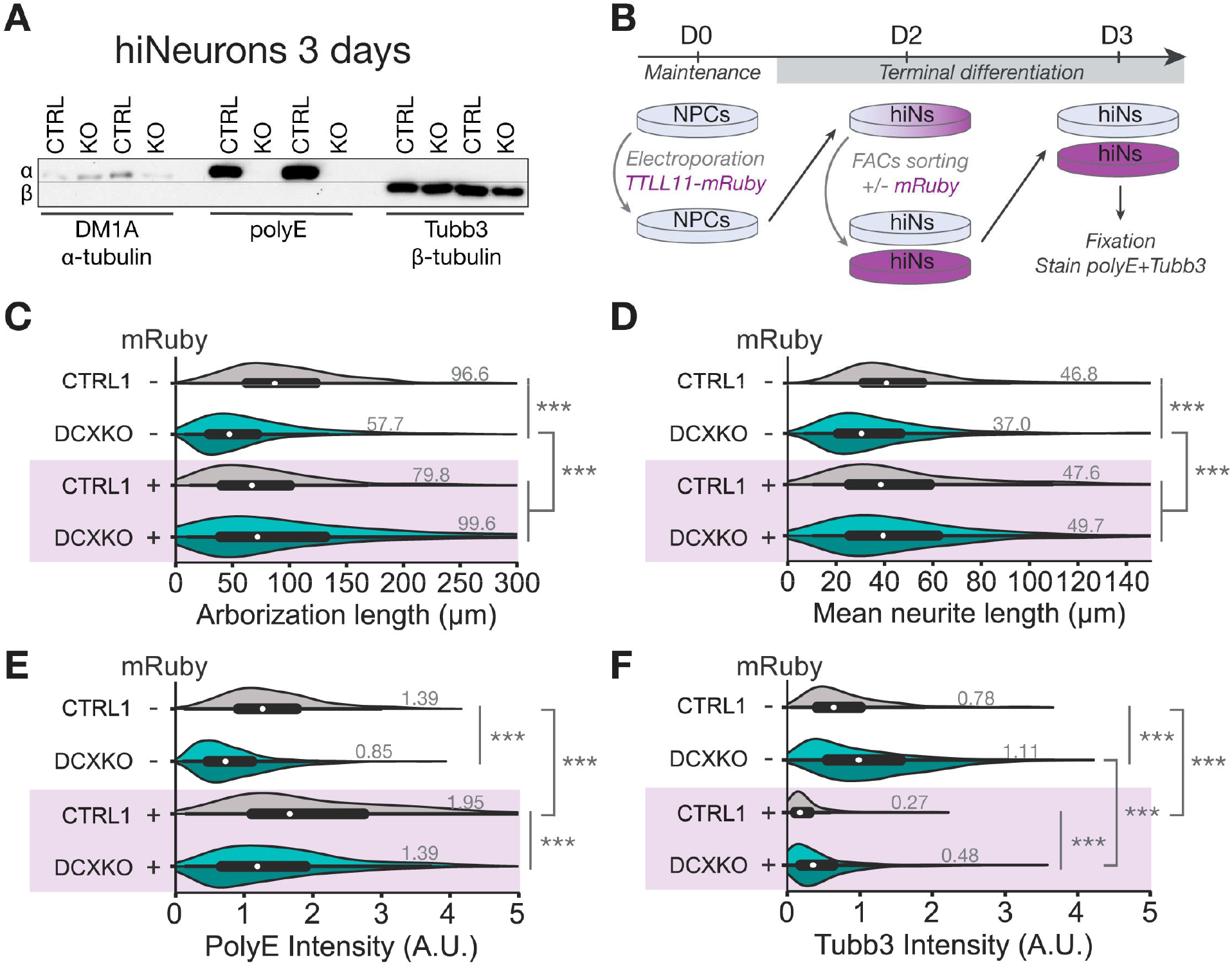
TTLL11 reduces neurite numbers by increasing tubulin polyglutamylation levels. **(A)** Immunoblots showing α-tubulin (DM1A) and β-tubulin (Tubb3) separately for CTRL and DCXKO (KO) DIV3 neurons. Polyglutamylation (PolyE) is visible only on α-tubulin subunits. Two sets of samples were loaded repetitively on the same gel and incubated with each antibody before re-alignement of the membranes for exposition. **(B)** Schematics of the experimental pipeline used for TTLL11 expression experiments. On D0, the NPCs were electroporated with TTLL11-mRuby and placed in terminal differentiation media. On D2 the cells were collected and sorted according to their mRuby signal (-/+). Once sorted the neurons (hiNs) were seeded at similar densities across conditions and left one more day in terminal differentiation media. On D3, the dishes were fixed and stained for polyE and Tubb3 as previously done. **(C)** Distribution of arborization length for fixed CTRL1 (dark grey), DCXKO1 (cyan) and DCXKO4 (dark cyan) neurons with or without TTLL11-mRuby expression. **(D)** Distribution of mean neurite length for fixed CTRL1 (dark grey), DCXKO1 (cyan) and DCXKO4 (dark cyan) neurons with or without TTLL11-mRuby expression. **(E)** Violin plots showing the polyglutamylation (polyE) intensity in CTRL1 and DCXKO DIV3 neurons with or without TTLL11-mRuby expression (mRuby -/+). **(F)** Violin plots showing the Tubb3 intensity in CTRL1 and DCXKO DIV3 neurons with or without TTLL11-mRuby expression (mRuby -/+). For C-F, images included in the analysis are from n=372 CTRL1, 380 DCXKO1, 654 DCXKO4, 199 CTRL1+TTLL11-mRuby, 186 DCXKO1+TTLL11-mRuby and 408 DCXKO4+TTLL11-mRuby neurons from 6 independent experiments. Statistical analysis was performed using a Mann-Whitney U test: *** *p*<0.001.

**Figure S11:**
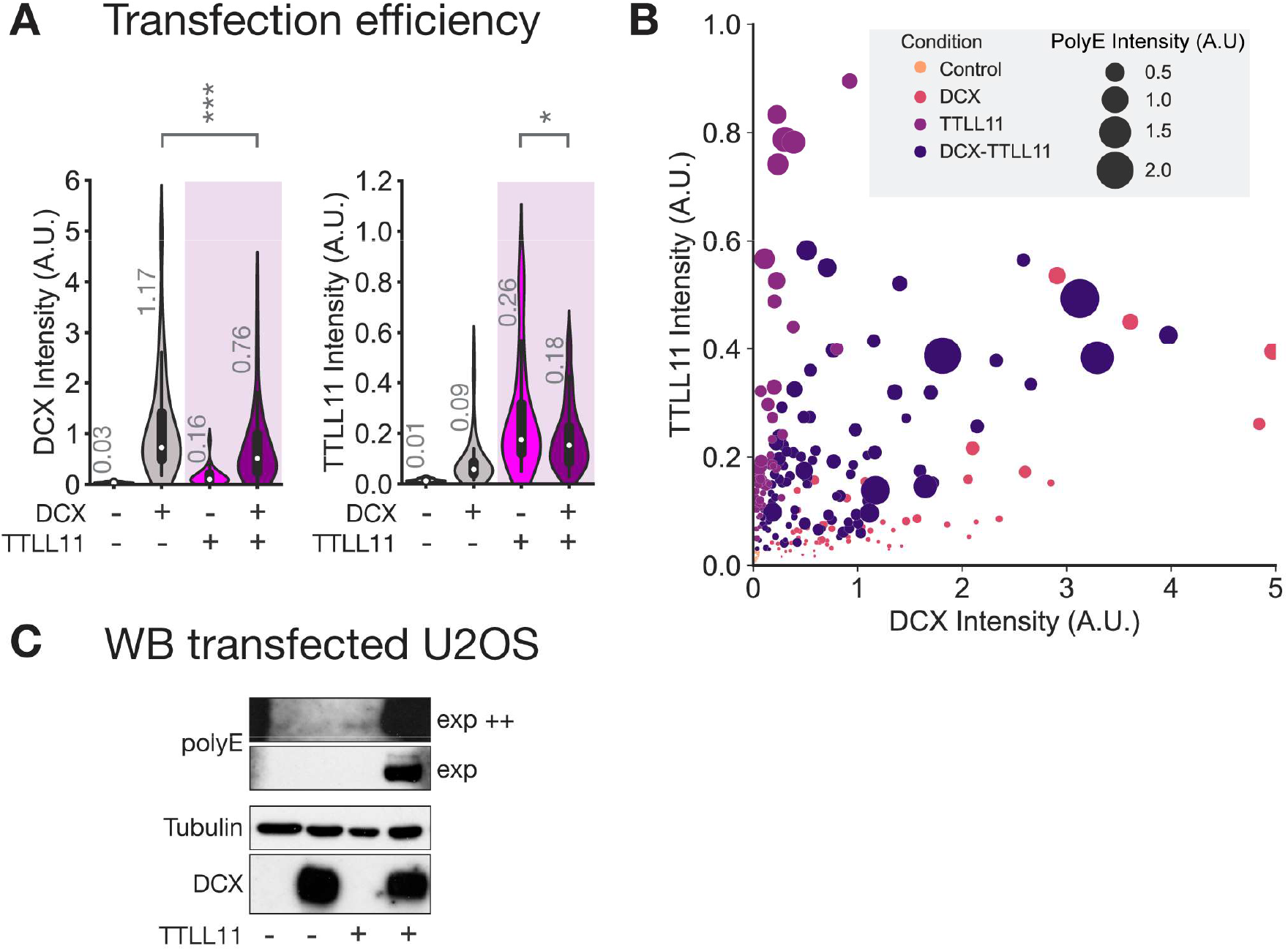
DCX and TTLL11 synergize to regulate polyglutamylation levels. **(A)** Violin plots showing the quantification of DCX-GFP (left panel) and TTLL11-mRuby (right panel) levels in immunostained U2OS cells. Images included in the analysis are from n=27 control (untransfected), 58 DCX-GFP, 46 TTLL11-mRuby and 95 DCX+TTLL11 cells from 3 experiments. Statistical analysis was performed using the Mann-Whitney non parametric U test: * *p*<0.05, *** *p*<0.001. **(B)** Scatterplot showing the U2OS production of polyglutamylation (polyE) depending on transfection conditions (untransfected or control, DCX-GFP, TTLL11-mRuby or DCX+TTLL11) and DCX-GFP and TTLL11-mRuby intensity. **(C)** Representative immunoblot showing polyglutamylation levels with two exposure time in U2OS cells transfected with DCX and TTLL11. TTLL11 expression was checked visually before sample collection. A small amount of polyE is visible in DCX and TTLL11 conditions when exposed for a long time (top panel).

