## Supplementary figures and images for "Doublecortin restricts neuronal branching by regulating tubulin polyglutamylation"

### Video S3_CTRL0 SiRTubulin CellMaskActin

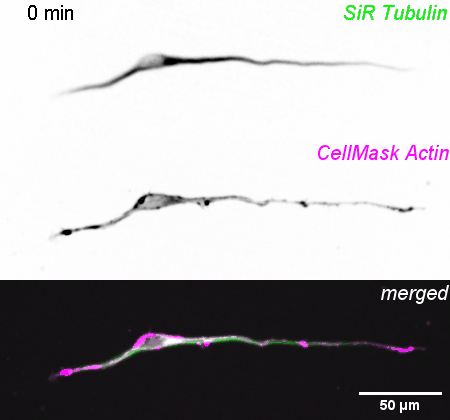

### Video S4_CTRL0 SiRTubulin CellMaskActin

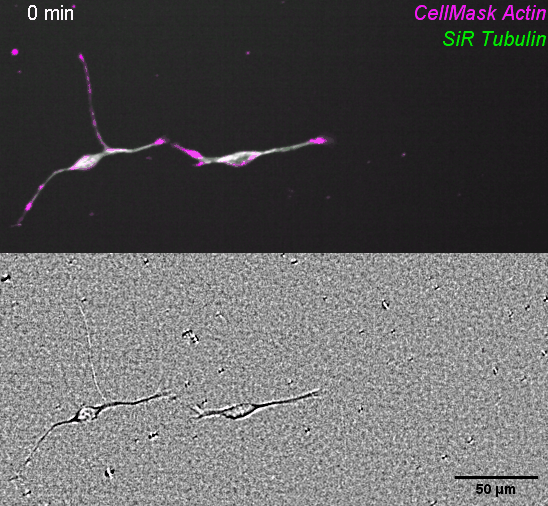

### Video S5_CTRL0 SiRTubulin CellMaskActin

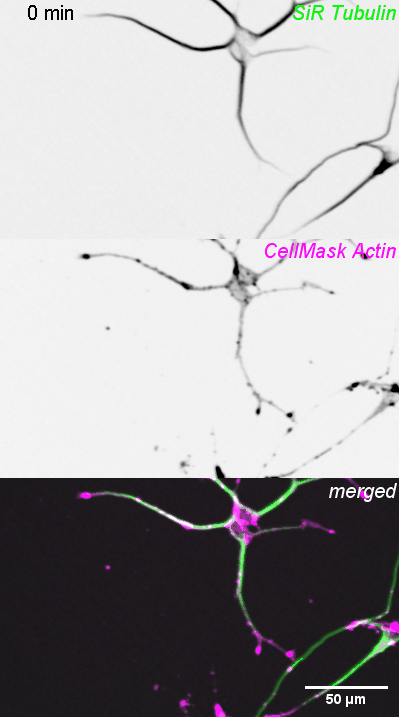

### Video S6_DCXKO1 SiRTubulin CellMaskActin

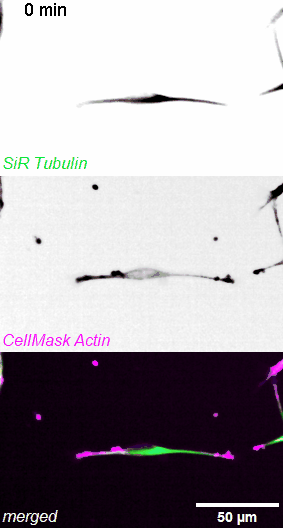

### Video S7_CTRL0 EB dynamics

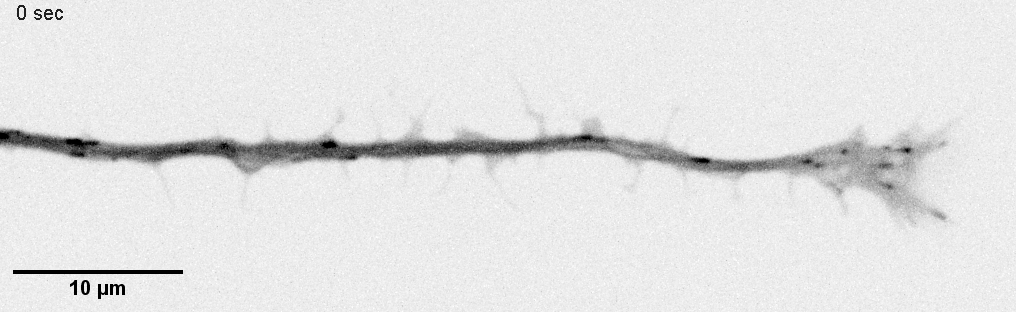

### Video S8_DCXKO1 EB dynamics

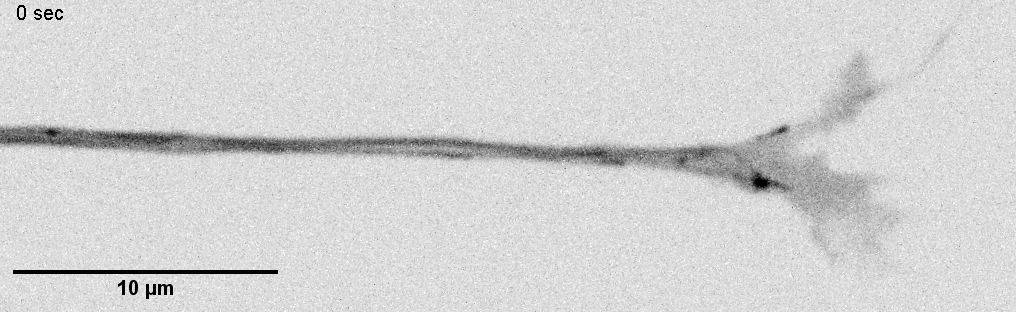

### Video S9_CTRL0 Lysotracker

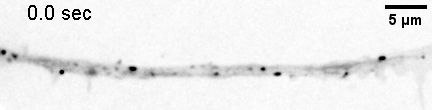

### Video S10_DCXKO1 Lysotracker

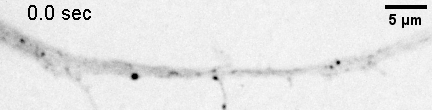
